# A glucanotransferase that uses two sub-sites and four catalytic aspartates/glutamates to disproportionate oligosaccharides ranging in length from maltotriose to starch

**DOI:** 10.1101/2023.08.25.554913

**Authors:** Arpita Sarkar, Pallavi Kaila, Prince Tiwari, Purnananda Guptasarma

## Abstract

PF0272 (PfuAmyGT) from *Pyrococcus furiosus* is a 656 residues-long, homodimeric, three-domain GH57 glycoside hydrolase [homologous to TLGT from *Thermococcus litoralis* (PDB ID: 1K1X)]. It is proposed to be either an α-amylase (EC 3.2.1.1), or a 4-α-glucanotransferase (EC 2.4.1.25). We demonstrate that PfuAmyGT is an exo-amylase-cum-glucanotransferase capable of transferring glucose, and dis-proportionating oligosaccharides, by excising glucose from malto-oligosaccharides (ranging in length from maltotriose to amylose/starch), and transferring it to malto-oligosaccharides (ranging in length from glucose to maltoheptaose and, possibly, even longer lengths). Convention holds that glucanotransferases transfer sugars through the serial and alternating binding of donors and acceptors to the same site, with covalent retention of excised sugars between such bindings. We present evidence of multiple behaviors in PfuAmyGT that challenge this view: (i) Production of free glucose, indicating scope for release of excised glucose; (ii) Higher activity with longer donors, indicating processivity; (iii) Accelerated activity with shorter acceptors, indicating a dependence upon rapid acceptor turnover; and (iv) Evidence of four catalytic glutamates/aspartates (E131, D222, E224, D362), indicating possible separation of excision and ligation functions. These behaviours collectively indicate the binding of donors and acceptors to separate sub-sites that have different substrate ‘length’ preferences, supporting our own previous proposal regarding separate tunnel (internal) and groove (surface) binding sites. Although PfuAmyGT’s mechanism of function remains to be fully elucidated, this paper definitively demonstrate ‘coupling’ of exo-amylase and glucanotransferase functions involving separate sub-sites for donor and acceptor binding.

## Introduction

Enzymes capable of modifying starch are of interest to enzyme biochemists, who are invested in understanding how such enzymes function, and also of interest to biotechnologists, who are invested in exploiting such enzymes towards specific applications.^1–3^ Thus, the search continues for new enzymes that are capable of degrading starch-based substrates, adding sugars to biopolymers, creating new sweeteners or anti-staling agents, and creating new edible gels from starch-derived saccharides.^4,5^

Glycoside hydrolase (GH) enzymes are sometimes also referred to as glycosidases or glycosyl hydrolases. GH enzymes are capable of hydrolysing glycosidic bonds, or joining carbohydrate moieties to other carbohydrate or non-carbohydrate moieties. About ∼180 GH families of enzymes are currently known [EC 2.4.1.X, 3.2.1.X, 4.2.2.X etc. (http://www.cazy.org/)].^6^ These include the two families that are discussed below, i.e., (i) GH13, which contains the well-known alpha-amylase enzyme, and (ii) GH57, which contains several atypical heat-stable hydrolases from *Dictyoglomus thermophilus*,^7,8^ and *Pyrococcus furiosus*.^9,10^ GH13 was established during the initial classification of glycoside hydrolases into different families.^11^ GH57 was established more recently, in 1996, however, as an afterthought.^12^

The *P. furiosus* enzyme known as PF0272 is considered to be a founding member of the GH57 family.^9,10^ Historically speaking, however, Anfinsen and colleagues had originally asserted, in 1993, that PF0272 is an alpha (endo) amylase, or GH13 enzyme, capable of hydrolysing starch.^10^ This assertion resulted in the use of the name, *amyA*, for the gene encoding PF0272 in the *P. furiosus* genome. However, within a few years, PF0272 was classified under GH57. Subsequently, Adams and coworkers,^13^ and also Park and coworkers,^14^ asserted that PF0272 is a dis-proportionating glucanotransferase. Adams and colleagues showed that the enzyme converts malto-oligosaccharides into pools of variously-sized oligosaccharides, maltose and glucose.^13^ Park and colleagues confirmed that PF0272 acts upon malto-oligosaccharides.^14^ Both groups demonstrated that the growth of *P. furiosus* upon maltose causes an up-regulation in the expression of PF0272. Such findings led to PF0272 being labelled a 4-α-glucanotransferase. Interestingly, neither Adams and coworkers^13^, nor Park and coworkers^14^, ever confirmed the presence of amylase activity in PF0272. On the contrary, Adams and coworkers actually explicitly asserted that PF0272 does not appear to catalyse starch degradation.^13^

GH57 family members typically contain five sequence motifs that are conserved in all homologous glucanotransferases of thermophilic/hyperthermophilic origin sequenced till date.^15,16^ According to some authors, 4-α-glucanotransferases (also known as a 1,4-α glucanotransferases) may be classified into five ‘types’ based upon the smallest donor and acceptor saccharides used by them, and also upon the glucan unit that happens to be transferred from the donor to the acceptor.^17,18^ According to this scheme of classification, PF0272 is thought to be a 4-α-glucanotransferase of Type V, i.e., an enzyme that uses maltose as its smallest donor, glucose as its smallest acceptor, and glucose as the sugar unit that is transferred.^18^

Our interest in PF0272, as an enzyme, was initially born out of a fundamental interest in understanding how a single enzyme could have been reported by two different sets of groups to be displaying functions as different as e.g., (i) an α-amylase (EC 3.2.1.1), and (ii) a 4-α-glucanotransferase (EC 2.4.1.25). Since this issue has remained unresolved ever since the work of Adams and coworkers,^13^ and Park and coworkers,^14^ happened to cast doubts upon the ability of PF0272 to process starch, we decided to clone, produce, and study PF0272 ourselves. Some years ago, in a preliminary paper describing our studies of this enzyme, we referred to PF0272 as PfuAmyGT,^19^ with ‘Amy’ referring to a potential amylase function, and ‘GT’ signifying a glucanotransferase function. Based on bioinformatics-based analyses and some preliminary experiments, we proposed a model for how the existence of two saccharide-binding sub-sites (instead of a single site) could allow PF0272/PfuAmyGT to function as both an exo-amylase, and a glucanotransferase.^19^

In the present paper, we demonstrate that PfuAmyGT excises glucose from donor saccharides to either release it (like an exo-amylase), or transfer it to an acceptor (like a glucanotransferase), with the efficiencies of these processes being clearly determined by the lengths and/or availabilities of donor and acceptor saccharides. We show that the PfuAmyGT’s smallest malto-glucan donor is maltotriose, and not maltose, as earlier assumed.^18^ We also demonstrate incontrovertibly that PfuAmyGT contains four separate aspartate/glutamate residues [E131 (domain 1); D222 (domain 1); E224 (domain 1); D362 (domain 2)] that function as either proton donor, or nucleophile base. The discovery of these four residues simultaneously confirms (i) our previous assertion that a loop residue in domain 2 (D362) is catalytically important,^19^ and also confirms (ii) a previous demonstration involving TLGT, the homolog of PfuAmyGT from *T. litoralis*, in which the residue, E123, in domain 1 (analogous to residue, E131, in domain 1 of PfuAmyGT) was shown to be catalytically important.^20,21^ Notably, in support of donors and acceptors binding to separate sub-sites, we had also previously referred,^19^ to the possibility of PfuAmyGT’s possessing at the interface of domains 2 and 3, a sugar-binding groove constituting what is often nowadays referred to as a surface binding site (SBS).^22,23^ We had proposed that this SBS could function as a second site in PfuAmyGT, in addition to a first site consisting of a tunnel in the enzyme leading to E131 (in domain 1) and D362 (in domain 2).

Viewing all of the available data and assessments of the enzyme, we propose that PfuAmyGT possesses (a) a processive donor binding sub-site located inside a tunnel, which prefers to act upon long malto-glucan chains (e.g., starch) in a processive manner, without releasing bound chains until they are shortened down to the length of two glucose units, i.e., maltose, and (b) an acceptor binding sub-site on the enzyme’s surface, i.e., an SBS that prefers to bind to shorter malto-glucans (e.g., maltose, maltotriose, maltotetraose etc.) which are rapidly turned over, with each glucan leaving immediately after the addition of a glucose unit. We propose that these two sub-sites are collectively serviced by four aspartate/glutamate residues [E131, D222, E224 and D262], and aided and regulated by concerted intra-domain loop motions, inter-domain motions, and inter-subunit motions. In a separate (companion) manuscript, we describe the construction and study of domain-swapped chimeras involving PfuAmyGT and another three-domain GH57 homolog called TonAmyGT (from *Thermococcus onnurenius*).^24^ TonAmyGT has recently been shown by us to use maltose (instead of maltotriose) as the smallest donor saccharide that it can work with,^24^ unlike PfuAmyGT which is shown to work with maltotriose as its smallest donor sachharide. In the companion manuscript, we show that chimeras incorporating domain 3 from PfuAmyGT happen to use maltotriose as smallest donor, whereas chimeras incorporating domain 3 from TonAmyGT use maltose as smallest donor. At present, domains 2 and 3 in GH57 enzymes are held to be domains of unknown function (DUFs).^25^ In this regard, both this manuscript and the companion manuscript suggest that domains 2 and 3 (and not just domain 1) play the following roles in the function of GH57 enzymes. We propose that a loop in domain 2 assists the transfer of glucose between two maltoglucan substrates that are bound simultaneously; one inside a tunnel in domain 1, and the other upon a surface binding site at the interface of domains 2 and 3. Clearly, more work remains to be done before the full mechanism of functioning of enzymes such as PfuAmyGT, TonAmyGT or TLGT (which we refer to TliAmyGT in the companion manuscript) become elucidated, or fully understood. Meanwhile, our two manuscripts provide significant scope for thought in regard of how conventional thinking about such glucanotransferases could need to be reviewed before GH57 enzymes can be fully understood.

## Materials and Methods

### Reagents

Starch and glucose were procured from HiMedia, India. Malto-oligosaccharide standards of different lengths were additionally procured from Santacruz Biotech or Merck (Sigma) USA. Isopropyl-1-thio-D-galactopyranoside (IPTG) was purchased from HiMedia or BR Biochem life sciences Pvt. Ltd, Delhi, India. For recombinant DNA work the restriction enzymes were procured from Fast Digest, Thermo Fisher Scientific and DNA polymerase and DNA ligase were purchased from New England Biolabs (MA, U.S.A.). TLC silica gel 60 F254 plates were procured from Merck, New Jersey, USA. Different kits like plasmid isolation kit, gel extraction kit and PCR clean-up kit was obtained from Qiagen, Hilden, Germany. For Site Overlapping PCR (SOE PCR), QuikChange® Site-Directed Mutagenesis Kit was procured from Stratagene, USA.

### Cloning and expression of PfuAmyGT and its mutants

The PfuAmyGT gene was re-amplified from a synthesized gene (optimized for expression in *E. coli*) and cloned in the pET-23a vector between *NdeI* and *XhoI* restriction sites. This plasmid was then inserted into XL-1 Blue *E. coli* cells. Subsequently, this vector was re-isolated and introduced into BL21 Star(DE3)pLysS cells for expression. The mutants in the present study were generated through use of SOE-PCR (as described earlier)^19^ or through use of the QuikChange® Site-Directed Mutagenesis Kit from Stratagene. Sequences of DNA primers used for PfuAmyGT and the point mutants are mentioned in Supplementary Information Table 1 (ST1).

**Table 1.** Calculated Cα-Cα Distance of selected residues (in Å) from D362 of the wild type PfuAmyGT.

| **Selected residues for mutation** | **Cα-Cα Distance from D362 (in Å)** |
| --- | --- |
| E32 | 19.1 |
| E33 | 18.8 |
| E131 | 9.5 |
| D222 | 14.4 |
| E224 | 15.2 |
| E623 | 16.4 |

### Heat Treatment and IMAC purification of PfuAmyGT

Transformed BL21 Star(DE3) pLysS cells expressing PfuAmyGT and the point mutants were used to set up primary culture containing appropriate antibiotics (100 µg/ml ampicillin and 35 µg/ml chloramphenicol) and incubated at 37 °C, with shaking at 220 rpm, for 12-16 h. This culture was used to inoculate 1 Litre of LB broth containing appropriate antibiotic and was allowed to grow until an O.D_600_ of 0.6, at which the inducer (IPTG) was added to a final concentration of 1 mM and the culture was allowed to grow further for 6-7 h at 37 °C or overnight at 25 °C. The cells were harvested through centrifugation at 7025 x g and re-suspended in non-denaturing lysis buffer (Qiagen) containing 10 mM Tris, 50 mM NaH_2_PO_4_ and 300 mM NaCl, with a pH of 8.0. The cells were lysed by sonication which was followed by heating the lysate at 90 °C for 15-20 minutes, to heat-aggregate the bulk of materials in the cytoplasm released through sonication. The heat aggregated lysate was subjected to centrifugation at 11290 x g for 1 hour to sediment the debris. The resulting supernatant, which was highly enriched in the protein, was passed through Ni-NTA resin (affinity chromatography), as per standard protocols (Qiagen). The eluted protein was dialyzed in 50 mM Tris buffer of pH 8.0 and electrophoretically examined on SDS PAGE using a Bio-Rad Tetrad apparatus, Richmond, CA, USA.

### Mass spectrometry-based protein identification

The identity of PfuAmyGT (77932.17 Da) was confirmed on a Synapt G2S-HDMS dual (MALDI/ESI) source Q-TOF LC-MS/MS mass spectrometer (Waters, USA), in three different types of mass spectrometric experiments: (i) intact protein mass determination, in the ESI mode, (ii) peptide mass fingerprinting (PMF), in the MALDI mode, and (iii) amino acid sequencing of a tryptic 1103 Da peptide, in the ESI mode. For accurate intact mass determination, the ESI source of the Synapt G2D-HDMS was employed, using the following experimental conditions: Positive ESI mode; direct infusion; m/z range, 50-5000; capillary voltage, 3 KV; cone voltage, 25 V; source temp, 100 ⁰C; desolvation temperature, 350 ⁰C; gas flow, 600 l/hr; sample flow, 50 µl/min. The raw ESI MS data was deconvoluted using the MaxEnt-1 function in the MassLynx software (Waters, USA). For peptide mass fingerprinting, and sequencing of the 1103 Da peptide, the MALDI source of the Synapt G2D-HDMS was used, under the following experimental conditions: m/z range, 50-5000; laser wavelength, 355 nm; sample matrix ratio, 1:1. The raw data was acquired using 100 ppm mass accuracy. PMF was performed by excising the ∼78 kDa SDS-PAGE band, subjecting it to in-gel alkylation/reduction and tryptic digestion, mixing it with the matric [α-cyano-4-hydroxycinnamic acid (CHCA)] in the ratio mentioned above, spotting it on the MALDI sample plate, drying it and subjecting to desorption by the mass spectrometer’s laser. Mass peaks obtained were compared to predicted masses from the sequence of PfuAmyGT, using the software tool, ExPasy. For peptide sequencing, the raw MALDI MS/MS data was deconvoluted using the MaxEnt-3 function of the MassLynx software, and the sequence was derived using the BioLynx tool of the MassLynx software. The obtained and expected amino acid sequences of the 1103 Da tryptic peptide were compared, for affirmation of the expected sequence.

### Determination of oligomeric status

The homodimeric structures of the purified PfuAmyGT and its point mutants were assessed through size exclusion chromatography (SEC), carried out on an AKTA Purifier 10 chromatographic system (GE Healthcare), using a Superdex-200 Increase 10/300 GL column, equilibrated with 50 mM Tris buffer [pH 8.0], at a flow rate of 0.5 ml/min. Eluted fractions were collected and analyzed on SDS-PAGE.

### Determination of secondary structural content

Circular Dichroism (CD) spectroscopy was performed on MOS 500 (Biologic) or Chirascan (Applied Photophysics) spectropolarimeters fitted with a peltier for temperature control. A quartz cuvette of 0.1 cm path length was used for collecting far UV CD spectra of protein samples between 200 and 250 nm. The mean residual ellipticity (MRE) was calculated in millidegrees using the equation, [Ѳ] = {Ѳ_obs_ (in mdeg) x 100 x MRW}/{1000 x protein concentration (in mg/ml) x path length (in cm)}, where the mean residue weight (MRW) was calculated by dividing the protein’s molecular weight (in Da) by the number of amino acids in its polypeptide chain.

### Confirmation of folding and presence of tertiary structure

The tertiary structures of PfuAmyGT and its mutants were checked by monitoring intrinsic tryptophan fluorescence for peak emission wavelength, on a Cary Eclipse fluorimeter, to examine whether the peak was at ∼353-355 nm (characteristic of tryptophan exposed to aqueous solvent) or below this wavelength (suggestive of tryptophan residues being buried through folding). Fluorescence emission spectra were recorded using excitation at 295 nm, and collecting emission spectra in the range of 300-400 nm. The slit width used was 5 nm for both excitation and emission monochromators, and the scan speed used was100 nm/min.

### Assessment of ultra-high thermal stability

Thermal denaturation data for PfuAmyGT and its point mutants was collected by monitoring changes in MRE at 222 nm as a function of temperature, through CD spectroscopy on a Chirascan (Applied Photophysics) spectropolarimeter fitted with a peltier for temperature control. Protein of 0.15 mg/ml concentration was taken in a sealed quartz cuvette of 0.1 cm path length, and temperature was raised from 20 °C to 90 °C, using 5 °C increments, at 5 minutes interval, with spectra collected at each temperature.

### Examination of activity. Starch-iodine method

Amylase activity was detected by the method of Fuwa,^26^ in which starch and iodine form a complex of blue-black colour, whereas oligosaccharides produced from hydrolysed starch react with iodine to form complexes of yellow colour. Starch (0.2 %) was mixed with 1 µM PfuAmyGT and incubated at 90 °C for 12 h (in a reaction volume of 20 µl). Following this, the mixture was mixed with 1 µl iodine (from a 1.25 % stock made in deionized water, and stored in the dark). This mixture was diluted appropriately, to measure A_620_ absorbance. A similar experiment was performed with 50 mM maltose to assess maltose-derived acceleration of PfuAmyGT’s activity upon starch at room temperature. *Zymogram method.* A suspension of 0.2 % starch with, or without, 200 mM maltose, was co-polymerized with 8 % acrylamide on a native PAGE, followed by the loading of 20 µl of PfuAmyGT (1, 2, 4 µM) onto the gel, prior to electrophoresis in the absence of SDS, and staining of the gel after electrophoresis by a 1 % iodine solution. Zones of clearance around the band corresponding to PfuAmyGT, caused by digestion of starch, were observed as readout. *Thin Layer Chromatography (TLC) method.* The amylase and glucanotransferase activity of PfuAmyGT and its mutants was assessed by TLCs, used to separate carbohydrates using highly polar absorbent silica as the stationary phase and a mixture of solvents as the mobile phase. In our case a mixture of Butanol:Ethanol:Water (50:30:20) was used. The oligosaccharides were separated on the basis of their relative polarities, through movement of the solvent front upon the TLC plate. The plates were dried, sprayed with a mixture of Methanol and Sulphuric acid in the ratio of 95:5, and heated at 120 °C for 2-5 minutes, to cause the charring/caramelization and visualization of sugar species.

### Determination of temperature, time and pH of optimum activity

The temperature of optimum activity was determined by TLC experiments (as described above) following incubation at different temperatures for 12 h. The time-dependence of PfuAmyGT activity was analysed by TLC, using aliquots from specific time points, through incubation at 90 °C, in reactions that were examined in respect of the differential abundances of various oligosaccharides generated. Likewise, the pH of optimum activity was also assessed through TLC, after incubation in different buffers of pH ranging from 2.0 to 12.0, at 90 °C for 12 h.

### Examination of acceleration of amylase activity by maltose

Under similar experimental conditions of incubation temperature and duration (as described above), acceleration of activity by maltose was checked by using a 1 % starch suspension in 50 mM Tris, pH 8.0, with an enzyme (PfuAmyGT) concentration of 1 µM (0.078 mg/ml), and using different concentrations of maltose (5-100 mM) and visualization using TLC.

### Mass spectrometry-based analysis of reaction products (malto-oligosaccharides)

After completion of the above enzymatic reaction (1 % starch + 20 mM maltose + 1 µM PfuAmyGT), oligo-saccharides obtained were diluted 100 times using mass spectrometry grade water, mixed in the ratio of 1:1 with the matrix, 2, 5-Dihydroxybenzoic acid [DHB; dissolved to a concentration of 10 g/L in 10 % (v/v) ethanol-water solution],^27^ using 1 µl each of the sugar hydrolysate and the DHB matrix. The mixture was spotted on MALDI plate(s) and subjected to mass spectrometry using the Synapt G2S-HDMS mass spectrometer (as already described earlier), to measure the masses of the sodiated ionic forms of different malto-oligosaccharides, using the same instrumental conditions earlier described for peptide mass fingerprinting.

### Identification of smallest donor and acceptor species

Experimental conditions were set up by using (i) glucose (20 mM) alone; (ii) glucose (20 mM) + PfuAmyGT (1 µM); (iii) maltose (20 mM) alone; (iv) maltose (20 mM) + PfuAmyGT (1 µM); (v) maltotriose (20 mM) alone; (vi) maltotriose (20 mM) + PfuAmyGT (1 µM). For all conditions above, experiments were carried out at two temperatures: 25 °C and 90 °C. Incubation was done for 12 hours. Products obtained were visualized using TLC.

### Processivity of PfuAmyGT

The processivity of PfuAmyGT was determined by comparing the activity of 1 µM PfuAmyGT (in the absence and presence of 5 mM maltose) upon 1.5 % solutions of starch, amylose and maltotriose (equivalent to 30 mM) incubated for 10 minutes at 90 °C and immediately analysed using visualization of products formed by TLC.

### Molecular visualization

The PyMOL Molecular Graphics System, Version 1.2r3pre, Schrödinger, LLC, n.d.,^28^ was used for visualizing protein structures, calculating intramolecular distances, and designing mutations. The pKa values of individual aspartate and glutamate residues were calculated using the Epik module of Schrödinger’s molecular modelling software.^29,30^

### Determination of protein concentration

Predictions of the extinction coefficients of PfuAmyGT, and its point mutants, at 280 nm, were made using the Expasy Protparam web server,^31^ by providing amino acid sequences (including information regarding numbers of different aromatic residues present). An optical density of 2.056 at 280 nm was predicted to correspond to a protein concentration of 1 mg/ml for PfuAmGT and all mutants.

## Results

### PfuAmyGT is an amylase that hydrolyses starch

The thin layer chromatogram (TLC) shown in Fig. 1A presents results of experiments examining whether PfuAmyGT acts upon starch. The experiment was conducted at two temperatures: (i) at room temperature (25 °C), and (ii) at a temperature (90 °C) commensurate with PfuAmyGT’s evolution in a hyperthermophile which is also feasibly accessed in a laboratory (without using the high pressure that ordinarily allows an archaeon like *P. furiosus* to grow at its usual growth temperature close to the boiling point of water). In lanes 1, and 7, respectively, controls corresponding to loadings of suspended starch (1 %) pre-incubated for 12 h at 25 °C, and 90 °C, are shown. In both cases, staining is seen in the thin layer chromatogram (TLC) only at the point (or spot) of loading, demonstrating the lack of any (detectable) enzyme-free hydrolysis of starch at either 25 °C, or 90 °C. In contrast, when PfuAmyGT (1 µM) is present in this suspension of starch, there are hints of the appearance of at least two malto-glucans (maltose and maltotriose) which begin to be seen, in lane 2, even for the incubation carried out at 25 °C for 12 h. For the incubation carried out at at 90 °C for 12 h, a substantially larger range of glucose and various malto-glucans are clearly seen in lane 8 of Fig. 1A (maltose, maltotriose, maltotetraose, maltopentaose, maltohexaose, maltoheptaose and longer species) with a much higher intensity of the spots corresponding to each oligo-saccharide species.

**Fig. 1.**
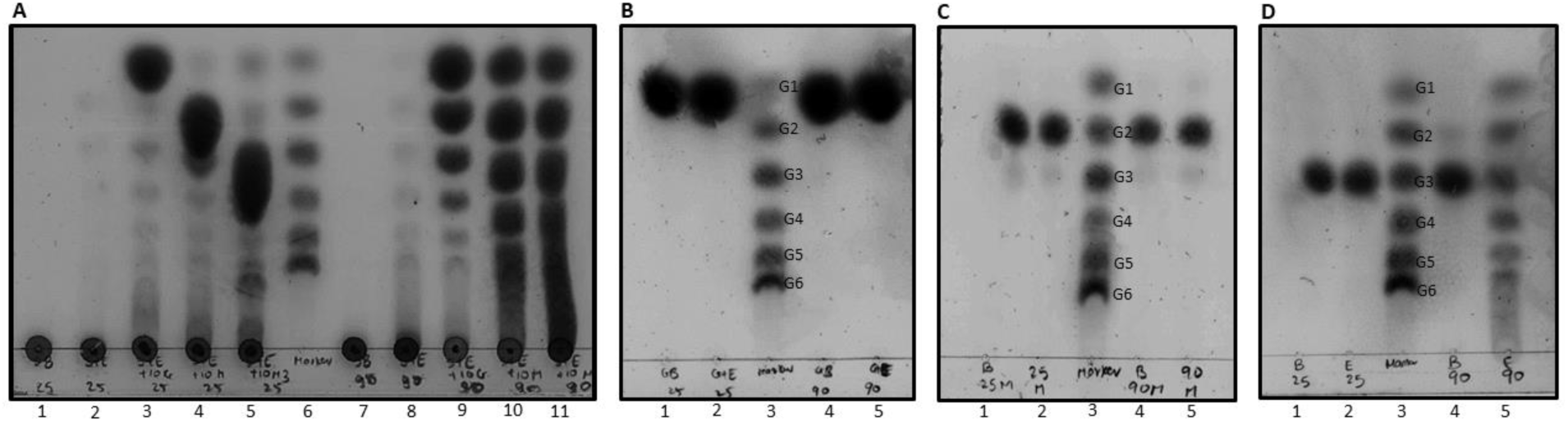
TLC experiments establishing the functioning of PfuAmyGT as both (i) exo-amylase, and (ii) a new type of maltose-accelerated (& temperature-accelerated) glucanotransferase using maltotriose as smallest donor, and glucose as both smallest acceptor & transferred glucan. **A**, TLC showing loading and mobility of starch (1%) following 12 h incubation with/without enzyme (PfuAmyGT, 1 µM) in the presence/absence of glucose/maltose/maltotriose (10 mM), at different temperatures. *Lanes 1 (25 °C) and 7 (90 °C)* correspond to starch in the absence of enzyme; *Lanes 2 (25 °C) and 8 (90 °C)* correspond to starch with enzyme; *Lane 3 (25 °C) and 9 (90 °C)* correspond to starch with enzyme and glucose; *Lanes 4 (25 °C) and 10 (90 °C)* correspond to starch with enzyme and maltose; *Lanes 5 (25 °C) and 11 (90 °C)* correspond to starch with enzyme and maltotriose; *Lane 6* shows premixed oligosaccharide markers containing 20 mM each of glucose (G1), maltose (G2), maltotriose (G3), maltotetraose (G4), maltopentaose (G5) and maltohexaose (G6). **B, C, D,** display TLCs showing the loading and mobility of 10 mM each of substrates such as glucose (panel b), maltose (panel c), and maltotriose (panel d), respectively, following 12 h incubation with/without enzyme (PfuAmyGT, 1 µM), at different temperatures, with no starch present. *Lane 1*: substrate with no enzyme at 25 °C; *Lane 2*: substrate with enzyme at 25 °C; *Lane 3*: premixed oligosaccharide markers identical to lane 6 of Fig.1a.; *Lane 4*: substrate with enzyme at 90 °C; *Lane 5*: substrate with enzyme at 90 °C.

We also examined PfuAmyGT’s ability to hydrolyze starch through use of the technique of zymography. The results of the zymography experiments are presented as side-by-side views of (i) Coomassie-stained native PAGE experiments in Supplementary Information Fig. S1, in which bands corresponding to dimeric PfuAmyGT (as well as a higher, multimeric form of PfuAmyGT) are seen, along with (ii) a starch-impregnated native PAGE that has been stained with iodine, which shows the same protein bands. Zones of clearance of starch are seen around the PfuAmyGT bands in the starch-impregnated gel, thus demonstrating PfuAmyGT’s ability to process/hydrolyse starch.

From all of the above data, it is clear that PfuAmyGT (PF0272) is capable of processing starch and turning it into smaller malto-glucans. It is also clear that PfuAmyGT hydrolyses starch much more efficiently at 90 °C, than at 25 °C, as would be expected for an enzyme evolved in a hyperthermophile archaeon. Our data here supports the earlier findings of Anfinsen and colleagues, who reported the ability of this enzyme to function an amylase,^10^ and questions the assertion made by Adams and colleagues,^13^ and also Park and colleagues,^14^ that PfuAmyGT does not act upon starch.

### PfuAmyGT’s amylase activity is accelerated by small saccharides like maltose

We next examined PfuAmyGT’s hydrolysis of starch in the presence of small malto-glucans, using experiments similar to the ones discussed immediately above. Fig. 1A shows results of experiments incorporating either glucose, or a small malto-glucan (such as maltose, or maltotriose) during incubations of PfuAmyGT with starch, both at 25 °C (lanes 3, 4 and 5), and also at 90 °C (lanes 9, 10 and 11). A significant increase can be seen in the production of glucose and other small glucans from PfuAmyGT’s processing of starch at 25 °C, in that oligosaccharides of multiple lengths are observed in lanes 3, 4, and 5, in comparison with lane 2 which shows only hints of maltose and maltotriose. At 90 °C, the presence of glucose and other small glucans causes a dramatic increase in the production of glucans of different sizes, as can be seen in lanes 9, 10 and 11. Further, at both 25 °C and 90 °C, the inclusion of maltose, or maltotriose, can be seen to elicit far more significant increases in the conversion of starch into different malto-glucans, than happens to be achieved through the addition of glucose. Although, of course, all three species accelerate the processing of starch, maltose is the most efficient in this regard than glucose, and maltotriose is more efficient than maltose.

We confirmed the above observations of an increase in starch hydrolysis by PfuAmyGT triggered by the presence of small glucans like maltose using one additional technique. The data for this is shown in Supplementary Information Fig. S2. Raw color photographs of residual starch stained by iodine, following enzymatic treatment at different temperatures, are shown in Fig. S2A. For the same samples, plots of the absorbance displayed by residual iodine-stained starch are shown in Fig. S2B. The panels in Fig. S2B clearly show that the processing of starch by PfuAmyGT is dramatically accelerated by the incorporation of maltose, at each of many different temperatures between 25 °C and 90 °C. Fig. S2C shows plate-based assays of starch clearance by PfuAmyGT, demonstrating larger zones of clearance for wells that additionally contained maltose, than for wells that contained only enzyme and starch, and no maltose. A simple comparison of lanes 4 and 10 in Fig. 1A make it evident that all the effects of the addition of maltose observed in the experiments presented in Supplementary Information Fig. S2 are derived from a dramatic acceleration in the action of PfuAmyGT to produce a range of maltoglucans of different sizes, whenever maltose is additionally present.

### The acceleration of PfuAmyGT’s amylase activity by maltose involves no allostery

Keeping in mind (i) the relatively-high maltose concentration (50 mM) demonstrated to exert activity-accelerating effects upon PfuAmyGT in Figs. S2A, S2B, and S2C, as well as (ii) the concentrations at which small molecules display allosteric effects, in general, which tend to be in the micromolar range when micromolar enzyme concentrations are used (and not in the range of tens, or hundreds, of millimolar maltose), it would appear that the activity-accelerating effects of maltose do not owe to any simple allosteric binding of maltose to PfuAmyGT. If the effect owed to simple allostery, no further effects of increasing maltose concentration would have been observed after the saturation of its binding site, whereas it is clearly seen in Fig. S2D that there is a progressive increase in the efficiency of processing of starch with increase of maltose concentration in the entire range from 5 mM to 100 mM, with no evidence of saturation, using a constant PfuAmyGT concentration of 1 µM. We have also examined this at 200 mM maltose (data not shown), i.e., a 2,00,000-fold excess of maltose over enzyme, with no signs of saturation.

Therefore, the acceleration of processing of starch by PfuAmyGT in the presence of maltose cannot be ascribed to any allosteric (high-affinity) binding of maltose to PfuAmyGT. Rather, as emphasized in subsequent sections, this progressive increase in yield of malto-glucans with progressive increase in maltose concentration results from maltose being used up as a substrate by PfuAmyGT, through the incorporation of maltose into various malto-glucan products that are made by PfuAmyGT from the excision and transfer of glucose from starch to maltose, and to malto-glucans of progressively longer lengths that are produced from maltose. In this regard, the following sections establish that PfuAmyGT is merely only incidentally an amylase. In the main, the enzyme is a 4-α-glucanotransferase (also known as a 1,4-α glucanotransferase, based on the type of glycosidic bond that is acted upon for glucan transfer, i.e., α-1,4 or 1,4-α glycosidic bonds).

Further, the amylase and glucanotransferase activities are also shown to be linked in a very interesting manner that has no precedent in the literature. In the sections below, we present data clearly indicating (a) that PfuAmyGT is a 4-α-glucanotransferase, and (b) also that its amylase activity (or ability to act upon starch) is actually an exo-amylase activity which functions in the service of this glucanotransferase activity, and which ‘feeds’ this glucanotransferase activity.

### PfuAmyGT is a 1,4-α glucanotransferase that dis-proportionates malto-oligosaccharides

The leitmotif of any 1,4-α glucanotransferase (or 4-α glucanotransferase) is that it transfers mono- or di-saccharides between a donor 1,4-α malto-oligosaccharide, and an acceptor 1,4-α malto-oligosaccharide). Further, the specific leitmotif of any dis-proportionating 1,4-α glucanotransferase is that it displays the ability to convert a single species of malto-oligosaccharide into a pool of malto-oligosaccharides of different lengths containing malto-oligosaccharides that are both shorter and longer than the original malto-oligosaccharide used as a substrate. This ability to both shorten and lengthen the original substrate, known as disproportionation, arises from the twin facts: (a) that, in each reaction cycle, the transfer of a mono- or di-saccharide shortens the donor oligosaccharide, and lengthens the acceptor oligosaccharide, and (b) that, between reaction cycles, both donor and acceptor oligosaccharides can exchange roles, such that over multiple reaction cycles the same oligosaccharide is sometimes shortened, and sometimes lengthened.

Figs. 1B, 1C and 1D, respectively, present the results of adding pure glucose, maltose or maltotriose to PfuAmyGT, in the complete absence of any starch, to examine whether any of these single (pure) sugars is transformed by PfuAmyGT into a pool of malto-oligosaccharides. Lanes 1, and 4, in each of Figs. 1B, 1C and 1D, respectively, present controls corresponding to experiments involving incubation of starch in the presence of glucose, maltose, and maltotriose, in the complete absence of PfuAmyGT, at 25 °C, and at 90 °C, respectively. Lanes 2, and 5, in each of Figs. 1B, 1C and 1D, respectively, correspond to experiments involving incubation of starch (1 %) in the presence of glucose, maltose, and maltotriose, in the presence of PfuAmyGT (1 µM), at 25 °C, and 90 °C, respectively. Fig. 1B shows that PfuAmyGT fails to lengthen glucose into maltose at 25 °C, and 90 °C, demonstrating that the enzyme cannot act as a glucanotransferase upon glucose alone, in the absence of starch or any other malto-oligosaccharide. Similarly, Fig 1C shows that PfuAmyGT fails to either shorten pure maltose into glucose, or lengthen it into maltotriose, at 25 °C, and 90 °C, demonstrating that the enzyme cannot act as a glucanotransferase upon maltose alone either, in the absence of starch or any other malto-oligosaccharide. In stark contrast, Fig. 1D shows that PfuAmyGT is able to both shorten maltotriose into maltose and glucose, as well as lengthen maltotriose into longer malto-oligosaccharides such as maltotetraose, maltopentaose, and maltohexaose, in the absence of starch or any other malto-oligosaccharide. Importantly, this ability of PfuAmyGT to process pure maltotriose is seen at 90 °C, but not at 25 °C. This may be contrasted with PfuAmyGT’s ability to act upon starch at 25 °C, when maltotriose (or maltose, or glucose) are present.

The above data clearly establishes that PfuAmyGT functions efficiently as a 1,4-α glucanotransferase upon small malto-oligosaccharides only at 90 °C, but not at 25 °C, although it is able to function efficiently as an amylase upon starch at both 90 °C, and 25 °C, in the presence of small malto-oligosaccharides that can presumably act as acceptors for glucose excised from starch through amylase activity. In other words, our results are made especially interesting by the facts that (a) although PfuAmyGT appears to be incapable of working with either pure maltotriose at 25 °C (Fig. 1D; lane 2), or with pure starch at 25 °C (Fig. 1A; lane 2), (b) it works extremely efficiently when both maltotriose, and starch, are present at 25 °C (Fig. 1A; lane 5). This evidence suggests that although maltotriose is the smallest donor with which PfuAmyGT is able to work as a 1,4-α glucanotransferase (Fig.1D; lane 5), starch turns out to be an even better donor than maltotriose, in the performance of this glucan transferring activity. This suggests that the site which is engaged in the binding of the donor saccharide displays some sort of a preference for starch over small glucans/sugars. Taken together with the other pieces of evidence demonstrating that these very same small malto-glucans, i.e., maltotriose (Fig. 1A; lane 5), maltose (Fig. 1A; lane 4), and glucose (Fig. 1A; lane 3) are all extremely efficient acceptors, unlike starch which is unable to act efficiently as an acceptor when starch itself is also the donor, as a malto-polysaccharide (Fig. 1A; lane 2), it appears to be clear that the site on the enzyme that is engaged in binding of saccharides functioning as acceptors displays a preference for small sugars over starch, whereas the site on the enzyme that is engaged in the binding of saccharides functioning as donors displays a preference for larger glucans like starch, over small sugars. In other words, PfuAmyGT prefers donor malto-oligosaccharides that are long, like starch, and acceptor malto-oligosaccharides that are short, like maltose, maltotriose et cetera. This very interesting aspect, which is extremely difficult to reconcile with the concept of a single site functioning to serially (and alternately) bind both donor and acceptor malto-oligosaccharides, is separately explored in greater detail, in a sub-section of the manuscript (after three intervening sub-sections that explore other aspects).

### Acceleration of starch hydrolysis by small saccharides indicates a coupling of exo-amylase function with glucanotransferase function

We draw attention here to the fact that precisely the same pool of malto-oligosaccharides, with apparently identical distributions of saccharide species of different lengths, happens to be generated by PfuAmyGT at 90 °C, through (a) action upon starch (alone), in the absence of any small malto-glucan (Fig. 1A; lane 7); (b) action upon starch in the presence of the small malto-glucan, maltotriose, (Fig. 1A; lane 11); and (c) action upon the small malto-glucan, maltotriose (alone), in the absence of any starch (Fig. 1D; lane 5). This suggests that the amylase and glucanotransferase functions of PfuAmyGT possess something in common that causes them to produce the same malto-oligosaccharide products. A possibility that springs to mind immediately is that these two functions of the enzyme may be coupled. In other words, PfuAmyGT could potentially be an exo-amylase that is capable of excising glucose from the ends of starch chains, or from the ends of small malto-oligosaccharides (with lower efficiency than that displayed with starch), with the ability of either (1) transferring this glucose to a waiting acceptor oligosaccharide, situated upon a different acceptor binding sub-site, or even (2) to an acceptor oligosaccharide that subsequently binds to the same sub-site, once it is vacated by the donor oligosaccharide. The latter possibility assumes that the donor oligosaccharide leaves the glucose covalently bound to a single sub-site at which it is later ‘picked up’ by the acceptor saccharide which binds to the same site. The former possibility, however, does not envisage such serial usage of the same site, but rather the simultaneous binding of donor and acceptor oligosaccharides to two separate but proximal sub-sites, with provisions for the transfer of an excised sugar between the donor and acceptor.

Notably, both of the above possibilities, between which we will distinguish later, are new possibilities that are neither in concert with the findings of Anfinsen and colleagues^10^ (who concluded that the enzyme is an endo-amylase that generates oligosaccharides of different sizes), nor in concert with the findings of the groups of both Adams,^13^ and Park,^14^ who concluded that the enzyme is not an endo-amylase or, indeed, an amylase of any sort or kind. The possibility that PfuAmyGT excises terminal units of glucose from starch and then either releases these into solution, or transfers them to a waiting (or subsequently-bound) acceptor potentially explains both (i) the generation of glucans of various sizes from starch, and (ii) the acceleration of starch-processing activity by the presence of small malto-glucans in the reaction, since the availability of acceptors apparently increases the rate of exo-amylolytic excision of glucose, by somehow mechanistically coupling such excision to the availability of an acceptor. The occasional miscarriage of such a coupling mechanism could potentially explain the increase seen in the rates of excision and release (or production) of free glucose, which is always seen. This production of free glucose is especially notable when a small malto-glucan is made available to PfuAmyGT as the sole substrate for processing as a 1,4-α glucanotransferase (Fig. 1A). Of course, at this point in the study, it is not clear why this should be the case. In subsequent sections, we show evidence supporting the possible existence of two separate (and non-overlapping) donor-binding and acceptor-binding sub-sites in PfuAmyGT.

### Maltotriose is established to be the smallest donor, while glucose is established to be both the smallest acceptor and the transferred unit

The fact that PfuAmyGT fails to lengthen glucose at 90 °C (Fig. 1B; lane 5), and also fails to either shorten or lengthen maltose 90 °C (Fig. 1C; lane 5), but is able to both shorten and lengthen maltotriose at 90 °C (Fig. 1D; lane 5), very firmly establishes maltotriose to be the smallest donor oligosaccharide that is used as a substrate, by PfuAmyGT. Of course, maltotriose is also one of PfuAmyGT’s acceptors at the same temperature (i.e., 90 °C), since maltotriose is not just shortened into shorter glucans, but also lengthened into longer glucans at this temperature. However, it is clear that maltotriose is not the smallest acceptor, since glucose clearly also increases the amounts of all species in the pool of malto-oligosaccharides produced by PfuAmyGT through hydrolysis of starch at 90 °C (Fig. 1A; lane 9). This clearly establishes glucose to be the smallest species that is used by PfuAmyGT as an acceptor, although it is clear that both maltose (Fig. 1A; lane 10), and maltotriose (Fig. 1A; lane 11) are more efficient acceptors than glucose.

Notably, some glucanotransferases are also thought to use maltose as the transferred unit. However, if maltose were the transferred unit in the case of PfuAmyGT, clearly the action of PfuAmyGT could only result in the visualization of glucose, maltotriose, and maltopentaose, following PfuAmyGT’s action upon maltotriose. In other words, there would be no production of maltose, maltotetraose, or maltohexaose, because transfer of maltose from one molecule of maltotriose to another molecule of maltotriose, or indeed to any further product generated initially from maltotriose (e.g., glucose, or maltopentaose), or generated subsequently (e.g., glucose, maltotriose, or maltopentaose) could only give rise to oligosaccharides containing odd numbers of glucose units. In contrast, what we observe clearly is the presence of oligosaccharides of every possible length ranging from glucose to maltohexaose, and varying by only single units of glucose (Fig. 1D; lane 5). Therefore, the transferred unit is clearly established to be glucose.

### PfuAmyGT appears to be a new type of 1,4-α glucanotransferase

We have already established that PfuAmyGT acts upon maltotriose to produce a pool of oligosaccharides, entirely independently of the presence of starch, with maltotriose initially acting as both donor and acceptor (Fig. 1D; lane 5), and also that maltotriose accepts glucose from starch (Fig. 1A; lane 11), just as maltose or glucose are able to do. On the other hand, we have also established that maltose generates a pool of malto-oligosaccharides by accepting glucose from starch, to turn itself into maltotriose, or maltotriose into maltotetraose, and maltotetraose into maltopentaose, and so on, into longer oligosaccharides (Fig. 1A; lane 10) but not when maltose is allowed to interacts with PfuAmyGT independently, in the absence of starch (Fig. 1C). According to the canon, no 1,4-α glucanotransferase is known to have these characteristics.^17,18^ Therefore, we suggest that PfuAmyGT could be a new type of 1,4-α glucanotransferase, i.e., it is certainly not the Type V enzyme that it has previously been assumed to be (based on classification by the use of substrates, and the misconception that it uses maltose as its smallest donor, rather than maltotriose).

### PfuAmyGT appears to use separate sub-sites for the binding and processing of donor and acceptor oligosaccharides

By convention, a glucanotransferase is assumed to possess a single site for the binding of donor and acceptor oligosaccharide substrates, with donor and acceptor oligosaccharides imagined to bind serially, and in a mutually-exclusive and alternating fashion, to this site. The donor is imagined to bind to this site leave a sugar covalently-bound to the enzyme in the form of an acyl intermediate, as a residue. The acceptor is imagined to bind to the same site subsequently, with the enzyme then transferring the covalently-bound sugar excised from the donor, to the acceptor. Of course, this mechanism provides a suitable theoretical explanation for how disproportionation can cause both shortening and lengthening of a single (pure) malto-oligosaccharide, since donors and acceptors can exchange roles and act, instead, as acceptors and donors, respectively, in successive rounds of glucan transferring activity.

However, the conception of the transfer (to an acceptor substrate) of a covalently-bound, donor-derived sugar, using a single substrate-binding site leaves no scope available for the conception of the types of counter-intuitive behavior that we have discovered to be exhibited by PfuAmyGT, in this regard. We find that the donor-derived sugar postulated to remain bound to the enzyme as an acyl intermediate (in this case, glucose) is released into solution in amounts that are comparable to those of the other small sugars that are produced through glucan-transfer. We also find clear differences in efficiencies of usage of starch, and maltotriose, respectively, as donor substrates (as shown in a previous section). Thus, in this section, we describe experiments conducted to fully explore and confirm both the counter-intuitive release of glucose, and the observed preferential usage of starch as a donor substrate, and maltotriose as an acceptor substrate (over starch).

### Glucose is released by PfuAmyGT

Fig. 2A shows a time-course for the production of different malto-oligosaccharide species by PfuAmyGT, when it is allowed to use maltotriose as a substrate. We find that there is no detectable production of any shorter or longer malto-oligosaccharide species at time-points corresponding to 0 min (lane 1), 2 min (lane 2), 5 min (lane 3) or 10 min (lane 4). Only the maltotriose substrate added initially is seen in the above lanes, when PfuAmyGT (1 µM) is incubated with maltotriose (20 mM) at 90 °C. However, at the 30 min time-point (lane 5), two products, i.e., maltotetraose and maltose, can be seen to have been formed in detectable quantities. These are, of course, the first two anticipated products of any disproportionation reaction involving maltotriose, since the two species are shorter, and longer, respectively, than maltotriose by a single glucose unit. At the 45 min time-point (lane 7), these two products, i.e., maltotetraose and maltose are seen to be present in greater amounts. In addition, maltopentaose and some maltohexaose, are also seen, with some glucose also beginning to be visible. At the time-points corresponding to 60 min (lane 8), 90 min (lane 9), 120 min (lane 10) and 240 min (lane 11), the main detectable further increase seen is in the amounts of glucose that are produced by the reaction.

**Fig. 2.**
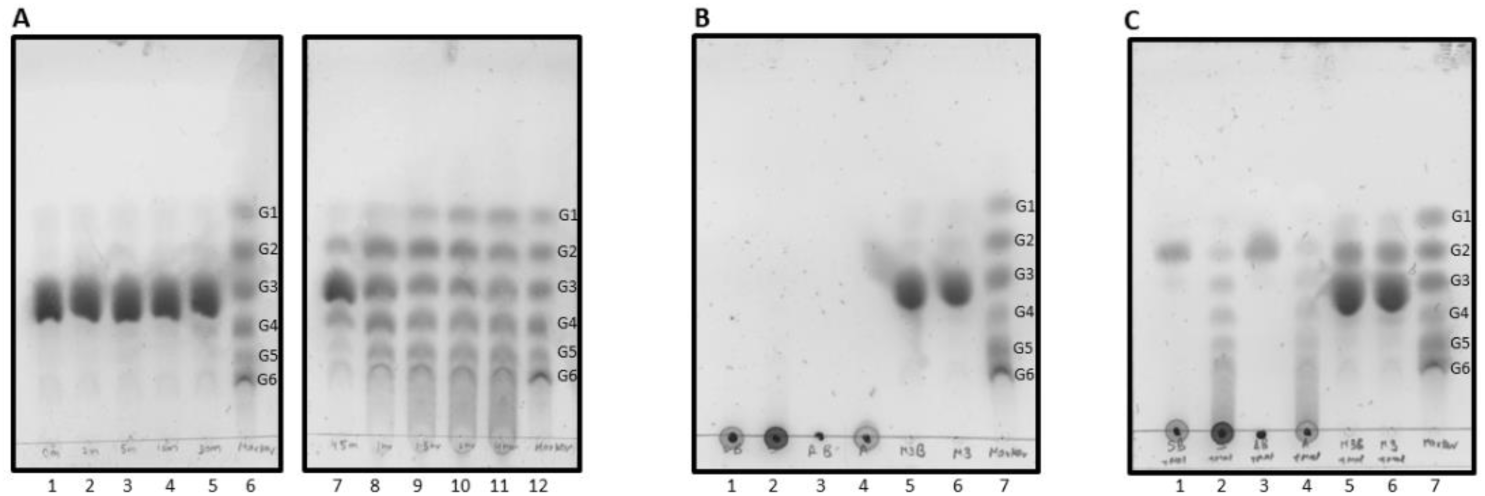
PfuAmyGT’s counter-intuitive production of free glucose and superior processing of starch/amylose over maltotriose. **A**, Time-course of the action of wild type PfuAmyGT upon 20 mM maltotriose. *Lane 1* shows the control containing maltotriose (20 mM) and PfuAmyGT (1 µM) frozen immediately after constitution. *Lanes 2, 3, 4, 5, 7, 8, 9, 10* and *11*, respectively, show products formed through 2, 5, 10, 30, 45, 60, 90, 120 and 240 min of incubation, at 90 °C, of 20 mM Maltotriose with 1 µM wild type PfuAmyGT. *Lanes 6* and *12* show pre-mixed standards (each 20 mM) containing varying numbers of glucose units, ranging to glucose (G1) to maltohexaose (G6). **B,** Examination of the relative amounts of detectable products formed, if any, through 10 min of incubation, at 90 °C, of equal w/v concentrations (1.5 %) of starch, amylose and maltotriose, with PfuAmyGT (1 µM). *Lanes 1, 3 and 5,* respectively, show enzyme-lacking controls of 1.5 % starch, amylose and maltose. *Lanes 2, 4 and 6,* respectively, show products formed through incubations of 1.5 % starch, amylose and maltotriose with PfuAmyGT (1 µM). *Lane 7* show pre-mixed standards (20 mM each), identical to lanes 6 and 12 of panel A. **C**, Examination of the relative amounts of detectable products formed during an experiment identical to that conducted and presented in panel B; the exception, in panel C, being that 5 mM maltose was added to all controls and all samples, to accelerate enzymatic reactions in samples shown in lanes 2, 4, and 6, presenting activity of PfuAmyGT upon starch, amylose and maltotriose, respectively.

The above data is indicative of two things: (i) *Attainment of equilibrium*: At some time-point between 120 min and 240 min, the intensities of bands corresponding to all six species, i.e., glucose, maltose, maltotriose, maltotetraose, maltopentaose and maltohexaose, become comparable between lanes in the TLC. This suggests that the disproportionation reaction has reached some sort of equilibrium, following which there is very little further difference in the distribution of products with further passage of time. In a subsequent section, we will note that a supplementary piece of data showing time-points that are even longer, i.e., up to 12 hours, serve to establish that this reaction does indeed attain equilibrium within about two hours. After this time-point, even if there is further enzyme-catalyzed interconversion of malto-oligosaccharide species through glucan transfer, clearly this occurs at similar rates for all species being transformed into other species, yielding no further changes in the distributions of amounts between species. (ii) *Release of glucose*: Until equilibrium is attained, glucose is very efficiently transferred, with almost no detectable build-up of glucose occurring that can be compared to the build-up of species such as maltose, maltotetraose, maltopentaose and maltohexaose, concomitantly with reduction in the amount of maltotriose (the originally supplied substrate). However, as equilibrium begins to be attained, and thereafter, there is very clear a progressive increase in the quantity of glucose until glucose too is in equilibrium with other produced species. This late-developing increase in the quantity of glucose establishes that the reaction does indeed produce glucose, as a sugar; not merely as a molecule that is transferred between substrates, but also as a molecule that is released into solution. Since glucose is also an acceptor substrate and not just the sugar that is transferred, presumably the transferred glucose is a part of the ‘donation’, and any released glucose is a part of the ‘acceptance’, while the reaction is running at fast rates, such that there is no build-up in the amounts of glucose. Once the rates of transfer slow down, however, as equilibrium is approached, scope presumably arises for the build-up of released glucose because it is no longer taken up by acceptor substrates at the same rate. Regardless of the correctness of the above explanation, we emphasize that the data clearly shows the production of glucose, which cannot be ignored under any circumstances. No theoretical means exist for PfuAmyGT to produce glucose, other than for the enzyme to occasionally release the glucose that is excised from the donor for the purposes of glucan transfer. This suggests that the enzyme has an exo-amylolytic function that is performed upon malto-oligosaccharide substrates that range in length from maltotriose to starch, with glucose being either releases, or transferred.

### Starch/amylose are superior to maltotriose as donors

A glucanotransferase with a single site that is serially, and alternately, occupied by donor and acceptor substrates in a mutually-exclusive fashion, may be anticipated to operate at comparable rates with all glucose-donating substrates (with especially long substrates potentially being processed at marginally slower rates, owing to diffusion-related issues). Therefore, if equal numbers of chain-ends are made available as substrates, an exo-amylase that happens to also function as a 4-α-glucanotransferase (e.g., an enzyme such as PfuAmyGT) could be expected to work at comparable rates when it uses any of the following substrates: (i) starch chains (containing mostly α-1,4 glycosidic bonds, and some α-1,6 glycosidic bonds), (ii) amylose chains (containing only α-1,4 glycosidic bonds), and (ii) maltotriose chains (containing only α-1,4 glycosidic bonds). However, if unequal numbers of chain-ends are made available, e.g., by providing equal w/v concentrations of starch, amylose, or maltotriose (ensuring identical numbers of glucose units and comparable numbers of glycosidic bonds), rather than equal molar concentrations (ensuring identical numbers of chain-ends), one anticipates that a small malto-oligosaccharide substrates (e.g., maltotriose) would produce the usual pool of malto-oligosaccharides more efficiently than starch or amylose, since it would be present in larger numbers for the same w/v concentrations, and also diffuse much faster both into and out of a single substrate binding site. Below, we show that experiments produce the exact opposite result which is commensurate not with a single site, but with processivity of use of starch/amylose and with separate donor and acceptor sites.

Fig. 2B and 2C show data for PfuAmyGT (1 µM) incubated with starch (1.5 %), amylose (1.5 %), and maltotriose (1.5 %; or 30 mM) for a period of 10 min at 90 °C. In each of Figs. 2B and 2C, lanes 1, 3 and 5, respectively, correspond to incubations of starch, amylose, and maltotriose in the absence of PfuAmyGT, whereas lanes 2, 4 and 6, respectively, correspond to incubations of starch, amylose, and maltotriose with PfuAmyGT. All lanes in Fig. 2B correspond to incubations performed without any added maltose, whereas all lanes in Fig. 2C correspond to incubations performed with maltose (5 mM) present. From lane 2, in Fig. 2B, it is clear that the enzyme is unable to produce detectable amounts of any of the species constituting the usual pool of malto-oligosaccharides, following a 10 min reaction with any of the supplied substrates. In contrast, when maltose is used to accelerate the reaction, as shown in Fig. 2C, clear distinctions become visible between the behavior of starch, or amylose, on the one hand, and maltotriose, on the other hand. Now, it is observed that PfuAmyGT produces copious amounts of all constituent species comprising the usual pool of malto-oligosaccharides, within a reaction of 10 min duration, when the enzyme acts upon either starch (lane 2), or amylose (lane 4), in the presence of maltose. However, when PfuAmyGT acts upon maltotriose (lane 6) in the present of maltose, over the same period of 10 min, only the reagents used (i.e., maltose and maltotriose) are visible, whereas none of the products comprising the usual pool of malto-oligosaccharides are visible. This demonstrates extremely clearly, and effectively, that PfuAmyGT processes starch more efficiently than it processes maltotriose, to produce the usual pool of malto-oligosaccharides, despite being present in smaller chain numbers than maltotriose, and also despite being bulkier than maltotriose and slower to diffuse. This demonstrates that the enzyme prefers starch as a donor over maltotriose, causing it to be at least as much of an amylase as a glucanotransferase. This also suggests that the site that binds to starch/amylose acts processively upon starch without releasing the starch chain between successive excisions of glucose from the end of the chain, since any release of starch between cycles of excision of glucose would cause it to be processed less efficiently than maltotriose, on account of its being bulkier and also present in smaller numbers, using comparable 1.5 % suspensions/solutions of both substrates.

It is worth noting that the result with maltotriose is consistent with data already presented in Fig. 2A, which describes experiments in which maltotriose was used at a somewhat lower concentration as a substrate (20 mM) than in the experiments described in Figs. 2B and 2C (30 mM). In Fig. 2A, we have already seen that the action of 1 µM PfuAmyGT upon 20 mM maltotriose does not begin to produce detectable species from the pool of oligosaccharides until nearly 45 min, with saturation observed between 120 and 240 min. Thus, it is not surprising that maltotriose does not give rise to detectable levels of the other species within 10 min, since these species only begin to become detectable by later time-points. What is very surprising, however, is that starch and amylose give rise to reasonably copious amounts of all species from the pool by the 10 min time-point. This point is so important that we wish to repeat it at the risk of being repetitive. Since we use the same w/v concentration of starch, amylose and maltotriose, this means that despite there being a lower number of chains of starch/amylose (both of which are much longer than maltotriose and, therefore, present in smaller numbers for similar w/v concentrations) and despite starch/amylose diffusing slower than maltotriose (due to their chains being longer), the pool of oligosaccharides is generated much earlier with starch/amylose than with maltotriose. This result cannot be explained if there is a single site that processes oligosaccharide chains in a serial fashion. However, it can be explained if there are two proximal sub-sites, one for donor binding and the other for acceptor binding, with the donor binding site functioning in a ‘processive’ manner. With such processivity involving the donor binding site, starch/amylose would not be required to dissociate from the donor binding site until the reduction of chains to the size of maltose (which cannot undergo further processing), explaining the more efficient processing of long substrates. Thus, the reduced requirement of donor chain dissociation/reassociation in comparison with maltotriose (reducible to maltose after the transfer of just one glucose unit) could very suitably explain the faster production of the pool of oligosaccharides from starch/amylose than from maltose.

In summary, the above data which demonstrates the faster processing of starch/amylose over maltotriose clearly indicates that a “two sub-site model” invoking separate and simultaneous binding of donor and acceptor explains the behavior of PfuAmyGT much better than a “one site model”, mostly because the former allows donor processivity to explain the faster processing of starch, whereas the latter offers neither scope for processivity nor any explanation for why longer chains should be processed so much more efficiently than short chains, despite being bulkier and present in lower amounts.

### Nominal confirmation of the identities of PfuAmyGT and its products

Before moving to mutational studies of PfuAmyGT to further understand its function, here we briefly describe results concerning the purification and confirmation of identity of PfuAmyGT, and of some of its products. *Purification and identification of PfuAmyGT.* As seen in lane 3 of Supplementary Information Fig. S3A, the SDS-PAGE gel band obtained following affinity purification of overexpressed PfuAmyGT from *E. coli* cell lysates is consistent with the predicted molecular weight of PfuAmyGT (∼78 kDa). As shown in lane 4 of Fig. S3A, the proteolytic degradation of PfuAmyGT seen in lane 3 is prevented by heating of bacterial lysates for 20 min at 90 °C, with this allowing the purification of PfuAmyGT to near homogeneity (ostensibly due to either heating-based inactivation of proteases, or heating-based facilitation of folding of this hyperthermophile-derived enzyme to completion, or both). Further purification through serial Ni-NTA affinity chromatography, and gel filtration chromatography, yielded an SDS-PAGE band for monomeric PfuAmyGT (Supplementary Information Fig. S3A) that was then subjected to (i) intact mass estimation, revealing a mass of 77953 Da against the expected mass of 77932 Da (Supplementary Information Fig. S3B); (ii) peptide mass fingerprinting (PMF) through in-gel tryptic digestion, to identify mass peaks matching peptide masses released upon trypsinization (Supplementary Information Fig. S3C, and Table ST2); and (iii) sequencing of a tryptic peptide (mass ∼1103 Da), yielding an amino acid sequence consistent with that of the said peptide [N-QQEIGEFPR-C], thus comprehensively establishing PfuAmyGT’s identity. *Identification of produced glucans.* Malto-oligosaccharides obtained through enzymatic hydrolysis of starch by PfuAmyGT in a manner accelerated by maltose were subjected to mass spectrometry (Supplementary Information Fig. S3D), leading to confirmation of mass peaks of 365.1023 Da (maltose), 527.1697 Da (maltotriose), 689.2419 Da (maltotetraose), 851.3149 Da (maltopentaose) and 1013.3824 (maltohexaose) Da, as sodium adducts of malto-glucans (Supplementary Table ST3).

### Both exo-amylase and glucanotransferase activities are simultaneously abolished by mutations, E131A, D222A, E224A and D362A, without detectable structural effects upon PfuAmyGT

We next explored the vicinity of residue D362, which had already been shown by us earlier to play a critical role in PfuAmyGT’s activity (please see the introduction section).^19^ This was done in order to identify D362’s partner aspartate/glutamate residue(s). Since D362 is located upon a loop in domain 2, and since this loop could be mobile [either autonomously, or in conjunction with movements of the other domains, as domain 2 happens to be sandwiched between domains 1 and 3 in a manner that could allow the other domains, and a loop upon domain 2, to potentially move back and forth between reaction cycles], we decided that we would examine the candidature of every single aspartate/glutamate residue lying within a C_α_-C_α_ distance of 20 Å from D362 within the structure of PfuAmyGT. The rationale for choosing a radius of 20 Å was that it is widely known that hydrolysis/formation of glycosidic bonds occurs through either a ‘retaining’ mechanism involving aspartate/glutamate residues separated by ∼5 Å, or through an ‘inverting’ mechanism involving aspartate/glutamate residues separated by ∼10 Å.^32,33^ Since the partner, or partners, of D362 could potentially lie to any side of D362, and since the distance separating the ends of an acarbose (maltotetraose analog) bound to domain 1 of PfuAmyGT and a maltose bound (unexplainedly) on the surface of domains and 3 in the crystal structure of a PfuAmyGT-homolog called TLGT,^20,21^ measures >15 Å, we arbitrarily chose a large sphere with a radius of 20 Å around residue D362, with a view to mutating every single aspartate/glutamate residue within this sphere to alanine, and examining the effects thereof upon PfuAmyGT’s activity.

The following six aspartate/glutamate residues were found to be located within a 20 Å radius of D362 in PfuAmyGT’s structure: E32, E33, E131, D222, E224, and E623 (see Table 1). We mutated these individually into alanine. Fig. 3A-3F present the results of our TLC-based examination of the abilities of these mutants to carry out (i) glucose excision and release, and (ii) glucose excision and transfer. Fig. 3A-3F show that the glucanotransferase activity of PfuAmyGT upon maltotriose (Fig. 1D; lane 5) was unaffected in three of the six mutants made: E32A (Fig. 3A), E33A (Fig. 3B), and E623A (Fig. 3F). In contrast, glucanotransferase activity was found to be severely affected in the other three mutants: E131A (Fig. 3C), D222A (Fig. 3D), and E224A (Fig. 3E), all three of which appear to have become incapable of carrying out any dis-proportionation of maltotriose, due to each of the three mutations, since no pools of oligosaccharides are observed to be generated. Further, we also reconfirmed (data not shown here) our earlier observation,^19^ that the fourth mutant, i.e., D362A, also completely lacks glucanotransferase activity. Therefore, including D362, a total of four acidic residues in PfuAmyGT appear to be critical for the enzyme to function as a glucanotransferase: E131, D222, E224 and D362.

**Fig. 3.**
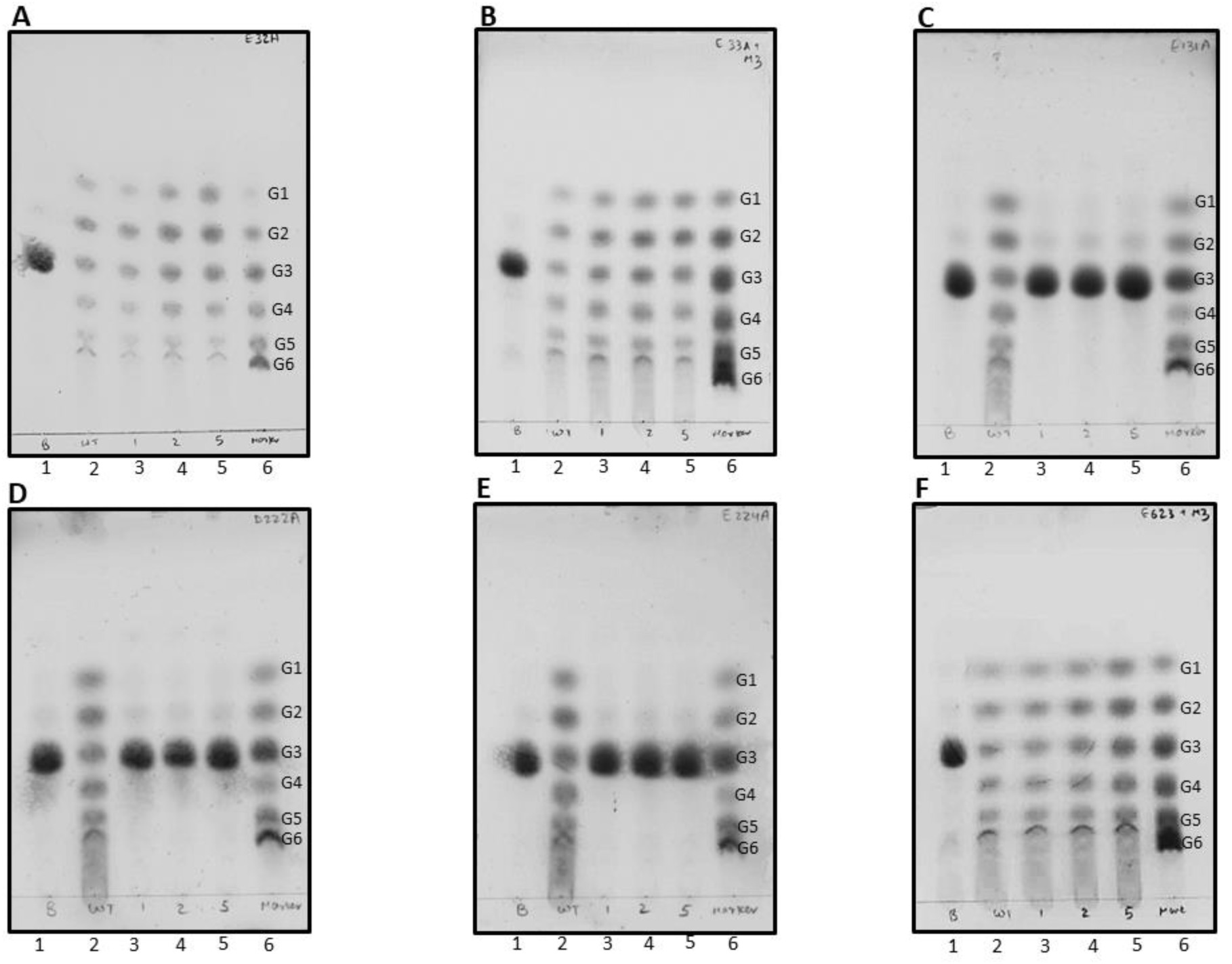
TLC experiments examining the retention/destruction of glucanotransferase function in 6 different mutants of PfuAmyGT. **A** (E32A)**, B** (E33A)**, C** (E131A)**,D** (D222A)**, E** (E224A)**, and F** (E623A), TLCs showing reaction products formed through 12 h incubations of enzyme (wild-type/mutant PfuAmyGT) with maltotriose (20 mM) at 90 °C. *Lane 1*: maltotriose with no enzyme; *Lane 2*: maltotriose with 1 µM wild-type enzyme; *Lane 3*: maltotriose with 1 µM mutant enzyme; *Lane 4*: maltotriose with 2 µM mutant enzyme; *Lane 5*: maltotriose with 5 µM mutant enzyme; *Lane 6*: premixed oligosaccharide markers containing 20 mM each of glucose (G1), maltose (G2), maltotriose (G3), maltotetraose (G4), maltopentaose (G5) and maltohexaose (G6).

### The enzymatically-inactive E131A, D222A, E224A and D362A variants are structurally indistinguishable from wild-type PfuAmyGT

To assess whether the observed mutation-induced loss of activity could derive from any significant change, or loss, involving the enzyme’s structure, we examined the following four types of behavior in each mutant of PfuAmyGT: (i) gel filtration chromatographic behavior (Fig. 4A), providing information about quaternary structure; (ii) intrinsic tryptophan burial-based fluorescence spectra (Fig. 4B), providing information about tertiary structure; (iii) circular dichroic spectra (Fig. 4C), providing information about secondary structure; and (iv) changes in thermal stability (Fig. 4D), providing information about more subtle changes that can increase the likelihood of chain unfolding. The above types of data was collected for all mutants displaying any loss of activity. Fig. 4A-4D show that the three activity-lacking mutants are similar to wild-type PfuAmyGT in every respect, establishing that loss of activity derives from the loss of catalytic ability. A similar examination of the other three mutants that remained active despite an introduced mutation is presented in Supplementary Information Fig. S4A, S4B, S4C and S4D. The relevant (similar) data for the D362 mutant has already been reported by us earlier.^19^

**Fig. 4.**
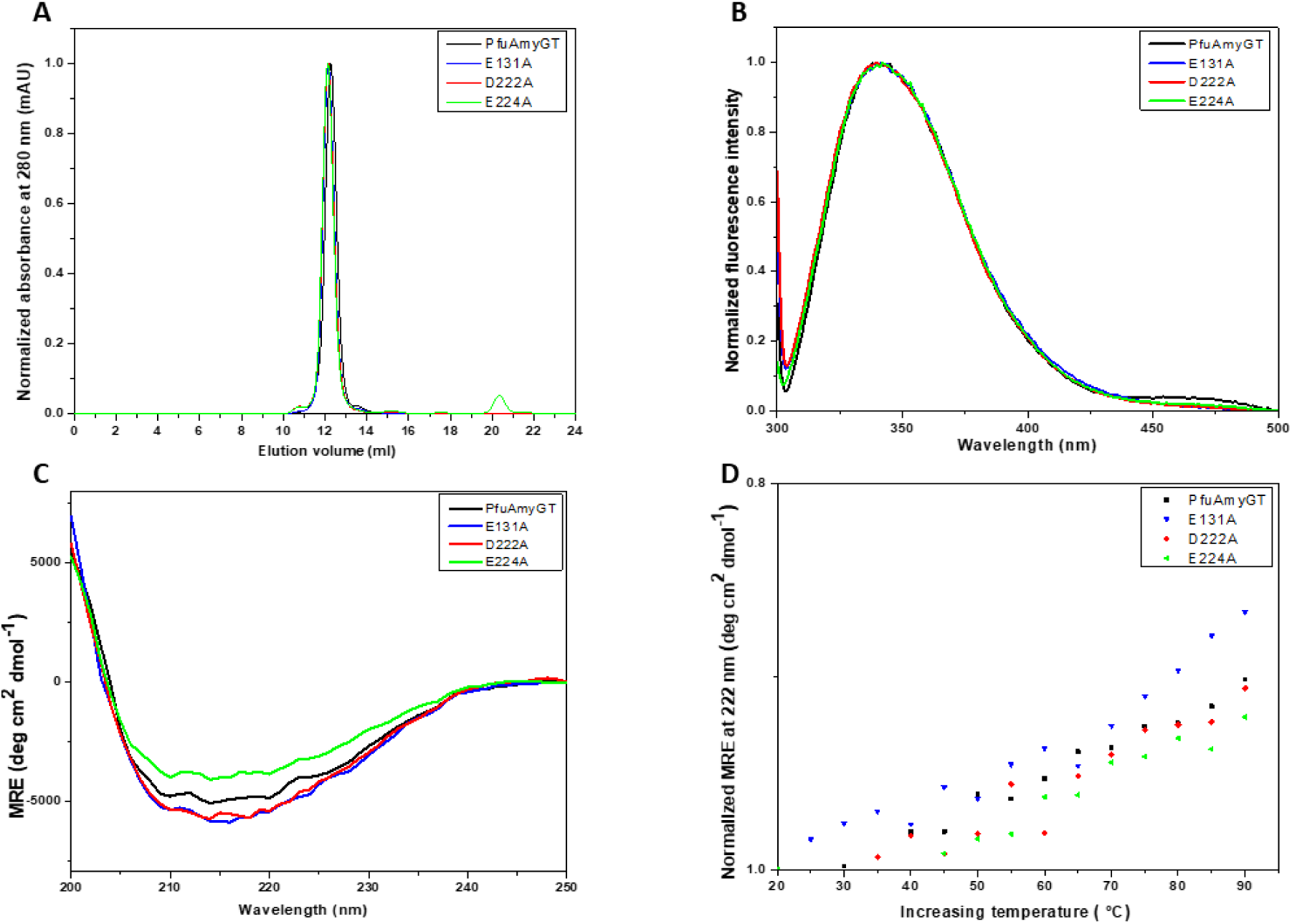
Comparative analyses of chromatographic and spectroscopic data for PfuAmyGT and its mutants, E131A, D222A, and E224A, in respect of the quaternary, tertiary and secondary structures of these proteins, and their thermal stability. **A** Gel filtration chromatographic profiles**, B** Normalized fluorescence emission spectra**, C** Circular Dichroism (CD) spectra**, D** CD signal intensity at 222 nm as a function of temperature.

### PfuAmyGT is folded and active over a wide range of temperature, time and pH

Supplementary Information Figure S5 shows PfuAmyGT’s activity as a function of varying temperature, time and pH. The data for temperature variations between 20 °C and 100 °C, in intervals of 10 °C are presented in Fig. S5A. The data for variations of time, varying from 2 h to 12 h, in intervals of 2 h are shown in Fig. S5B. Notably, the data for time-points less than 2 h have already been presented in Fig. 2A. The data for variations in pH, between pH 2.0 and pH 12.0 are presented in Fig. S5C. From the presented data, the following conclusions may be drawn. Firstly, we conclude that the enzyme is active in the presence of maltotriose over a wide range of temperatures between 60 °C and 100 °C, with peak activity seen at 90 °C. This is not surprising, considering the fact that the enzyme’s sequence, and three-dimensional structure, evolved inside the genome/proteome of a hyperthermophile organism like *P. furiosus*. Secondly, we conclude that the enzyme achieves equilibrium between oligosaccharide species within two hours of incubation, under the conditions tested, with little further change in the distribution of species at longer time points, supporting a conclusion reached in an earlier section. Thirdly, we conclude that apart from some lowering of activity in the vicinity of pH 7.0, for which we currently have no explanations, the enzyme appears to be active over a wide range of pH, between pH 3.0 and pH 12.0, with peak activity at pH 9.5, i.e., in a slightly alkaline range of pH.

### Prediction of the pKa values of E131, D222, E224 and D362, and implications of these values

We examined the pKa values of the four catalytically-important asparate/glutamate residues, as predicted by the Epik module of Schrödinger’s molecular modelling software.^29,30^ These are given below under parentheses for each of the four residues: E131 (pKa 7.68); D222 (pKa 4.71); E224 (pKa 2.73); D362 (pKa 4.95). Given its very low pKa, residue E224 (with a pKa of 2.73) is likely to be a nucleophile, acting with help from D222 and in concert with residue E131 to initiate the hydrolytic removal of glucose from donor saccharides. It is possible that the glucose is then either released or transferred to D362, with D362 being somehow tasked with carrying the glucose into the proximity of the acceptor, followed by the ligation of the glucose to the acceptor, in cooperation with either E131 or E224 which move into the vicinity through motions in the enzyme. It is also possible that intra-domain (loop) and/or inter-domain movements allow two of these residues to aid the hydrolytic removal of a glucose from the donor, and the other two residues to aid its ligation onto the acceptor, with assistance from W365. Notably, W365 shows an effect upon being mutated to alanine.^19^ The actual combinations in which these four residues contribute to what happens at the donor-binding and acceptor-binding sub-sites are likely to be influenced by movements in the enzyme, and it is difficult to propose a mechanism without understanding these motions. It is also conceivable that there is the formation of an intermediate covalent glycosyl ester during the transfer process.

## Discussion

Carbohydrate-processing enzymes host saccharide/glucan-binding sites that constitute any one of the following structural features upon their surfaces: (i) a pocket, or crater; (ii) a cleft, or groove; and (iii) a tunnel, or pore.^33^ It is commonly assumed that a single enzyme contains only one of these three features; however, this is not a dictum that prevents consideration of additional saccharide-binding sites. Indeed, in recent years, additional sites that are called surface binding sites (SBS) have been observed and commented upon,^22,23^ with some of these sites also appearing to influence substrate binding or specificity. Little is known about the functions of these surface binding sites. In this paper, we have imputed a role for a surface binding site at the interface of domains 2 and 3 (previously described by us to be a site that binds to maltose)^19^ as a sub-site of the active site which is involved in the binding of acceptor substrates. In a companion paper, we have imputed a role for this SBS in determining donor binding specificity at a proximal site within domain 1.

It is commonly assumed that a single enzyme contains only a single pair of catalytically-important acidic (aspartate/glutamate) residues, and that this pair is alternately involved in the hydrolysis of a glycosidic bond in a donor saccharide, and in the formation of a glycosidic bond in an acceptor saccharide, when the enzyme is a glucanotransferase. In other words, the same pair of residues is assumed to be involved in both the shortening of the donor, and in the lengthening of the acceptor, since the transferred sugar unit is assumed to remain covalently bound at a site which is serially, and alternately, occupied by the donor (which deposits the cleaved sugar, at the site) and the acceptor (to which the cleaved sugar become ligated, at the same site). However, in this paper, and in the case of PfuAmyGT, we have demonstrated several features of the enzyme that argue against the existence of both a single substrate-binding sub-site, and also against a single pair of catalytic aspartates/glutamates.

Briefly, we have demonstrated the following: (1) that the enzyme is both an amylase and a glucanotransferase; (2) that the amylase and glucanotransferase activities appear to be coupled; (3) that the amylase activity is an exo-amylase activity (and not an endo-amylase activity); (4) that the exo-amylase activity operates in the service of the glucanotransferase activity, and also under the regulation of the glucanotransferase activity, in that the availability of small malto-oligosaccharide acceptors (like maltose) very severely determine the rate of processing of long donor substrates such as starch, or amylose; (5) that the enzyme can both transfer glucose from a donor substrate to an acceptor substrate and also release this glucose into solution; (6) that the enzyme clearly prefers starch/amylose as donor, over small malto-oligosaccharides (like maltotriose); (7) that the enzyme appears to use a processive mode of action against starch that allows it to be progressively shortened, potentially without being released between successive excisions of glucose; (8) that the enzyme clearly prefers small malto-oligosaccharides (like maltotriose) as acceptor, over starch/amylose, with evidence for the rapid cycling/turnover of acceptors to produce a pool of oligosaccharides; (9) that the enzyme uses maltotriose as its smallest donor, glucose as its smallest acceptor, and glucose as the sugar that is excised and transferred, or released; (10) that the enzyme appears to use four catalytic glutamates/aspartates, E131, D222, E224 and D363, rather than just two catalytic glutamates/aspartates; (11) that D362 (in domain 2) is at a distance from E131 (in domain 1), suggesting the requirement for inter-domain motions during the catalytic cycle; (12) that D362 lies between E131, D222 and E224 located at the end of a tunnel in domain 1, and a groove-shaped, maltose-binding, SBS in domain 3, which is very suggestive of the existence of two separate sub-sites for donor and acceptor binding. Our preference is for the tunnel in domain 1 to act as the donor binding site, since a tunnel reduces the scope for rapid turnover of substrate and can facilitate processivity in the excisions of successive glucose units from the end of a long donor starch chain for transfer to surface-bound acceptors that are easily turned over, with the loop carrying D362 in domain 2 somehow facilitating the transfer.

The above discussion supports the proposal that PfuAmyGT, and other GH57 enzymes like it, have been designed by nature for cleaving a glucose from starch/amylose and transferring it to a glucose, or maltose, bound at a separate site, through the movement of some component of the enzyme. In a previous paper too, without all of the above evidence, but with some bioinformatics-based analyses of available crystallographic data,^21^ we had proposed that PfuAmyGT possesses two separate saccharide-binding sites.^19^ Briefly, we had pointed out that a non-hydrolysable analog of maltotetraose, known as acarbose (a known inhibitor of glucanotransferase function) was reported to be bound inside a tunnel upon TLGT, a *Thermococcus litoralis* glucanotransferase homolog of PfuAmyGT sharing 64.6 % sequence homology, following addition of acarbose during crystallization of TLGT. The structure of TLGT published by Imamura and coworkers, exists in the published literature (PDB ID: 1K1X).^21^ Like PfuAmyGT, which behaves like a dimer in gel filtration studies, the *T. litoralis* glucanotransferase is a homodimer in the crystal structure, with acarbose bound to a tunnel-like feature in only a single subunit, and not to both (identical) subunits, with the non-hydrolysable (nitrogen atom-containing) terminus of acarbose pointing into the tunnel’s recesses. We combined this information with the fact that the electron density of a maltose molecule (not added to the crystallization set up) was reported to be puzzlingly seen upon the other (acarbose-lacking) subunit of TLGT. We had thus speculated that maltose was picked up from the cytoplasm of the *E. coli* expression host of TLGT, and suggested that the presence of the maltose indicates a surface binding site for saccharides upon TLGT. We structurally transposed this maltose molecule onto the acarbose-bound subunit, and proceeded to map both acarbose and maltose onto a structural model of PfuAmyGT (built using homology-based modelling using TLGT as template), to show that (i) the acarbose is bound to a tunnel upon domain 1; (ii) that acarbose can potentially bind proximally to the transposed maltose, bound to a groove upon the enzyme’s surface (lined by residues Q576, S577, and Y596, from domain 3; residues H376, L377 and A380 from domain 2; and residue G28 from domain 1), and (iii) that the acarbose-bound tunnel and maltose-bound groove are separated by a loop from domain 2 (residues 360-374) containing residues D362, and W365, which abolish, and diminish, respectively, the activity of PfuAmyGT, upon mutation to alanine. The above findings had then caused us to propose that W365 helps in the transfer of glucose between donor and acceptor saccharides, and that D362 is a catalytically important aspartate/glutamate.^19^

In Fig. 5 below, we present a view of the regions of a single subunit of the modelled structure of PfuAmyGT (with an expanded view shown alongside), highlighting the parts of the molecule proposed to be involved in saccharide binding and catalysis. The maltose, and acarbose (maltotetraose analog) are shown in ochre, at the top left, and bottom right, respectively, of the expanded view, in Fig. 5. The aspartate residue already demonstrated to be catalytically important,^19^ i.e., residue D362, is shown in green. The aspartate/glutamate residues that are present within a 20 Å radius of D362 (E131, D222, E224, E32, E33, E623), and which were each individually mutated to alanine, for the experiments reported in this paper, have been shown in red. The figure makes it evident that the probability of involvement of the loop containing D362 in the transfer of glucose between the two saccharides is significantly high. It is also evident that the residues shown in this paper to be catalytically important, i.e., E131, D222 and E224 are close to the nitrogen atom of the acarbose which acts as a proxy for the oxygen in the glycosidic bond of maltotetraose.

**Fig. 5.**
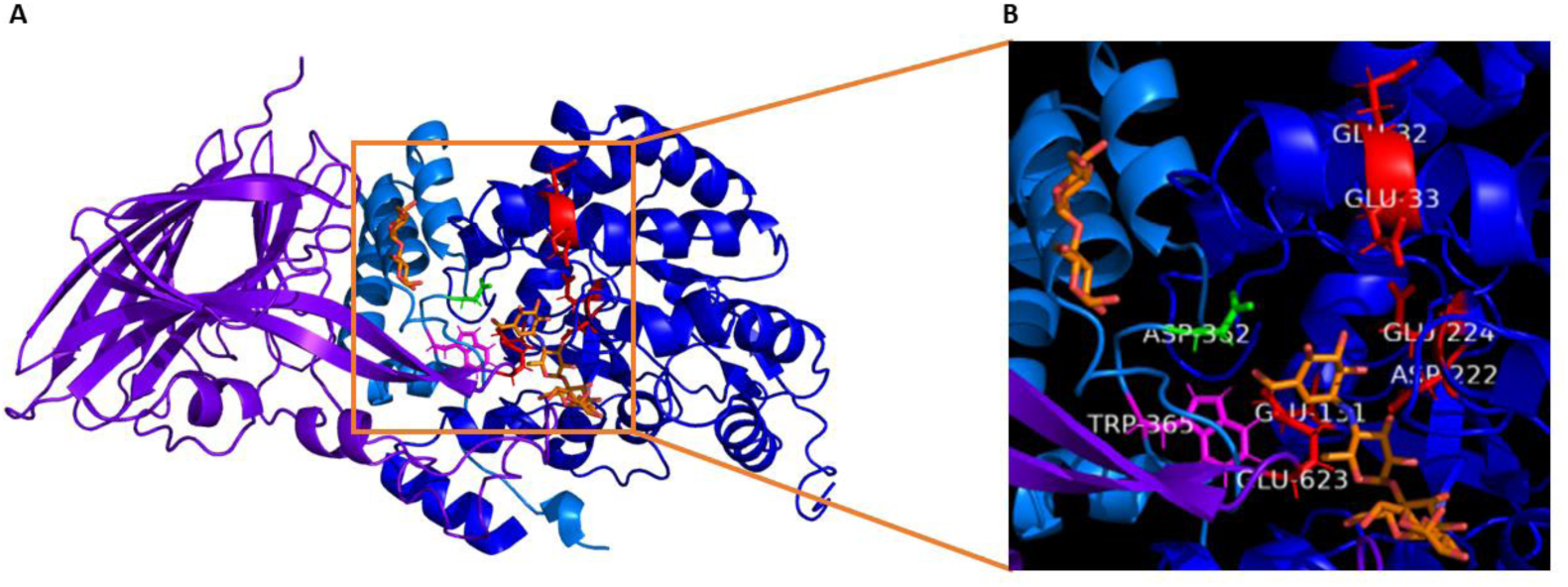
Views of PfuAmyGT and its proposed catalytic core. **A,** Homology modelling-based structure of PfuAmyGT (using a *T. litoralis* homolog as template) showing a single subunit of the enzyme homodimer, and domains 1 (blue), 2 (grey) and 3 (purple), with acarbose bound inside a tunnel in domain 1, and maltose shown bound to a surface groove consisting mainly of residues from domains 2 and 3. In the *T. litoralis* homolog, acarbose and maltose are bound mutually-exclusively to two different subunits. **B,** Expanded view of the figure shown in panel a, showing acarbose and maltose at the proposed catalytic site, highlighting the locations of seven residues E32 (red), E33 (red), E131 (red), D222 (red), E224 (red), D362 (green), and E623 (red). The residues in red lie within a 20 Å radius of the residue (D362) shown in green

Notably, Imamura and coworkers had speculated upon the residues that could constitute the canonical catalytic pair in the *T. litoralis* glucanotransferase.^20,21^ We find that these residues turn out to be analogous to two out of the four residues that we have established to be catalytically important, namely residues E123 in the *T. litoralis* enzyme (analogous to E131 in PfuAmyGT) and D214 in the *T. litoralis* enzyme (analogous to D222 in PfuAmyGT). On the other hand, residues analogous to D362 (which was the first residue to be confirmed by us to be catalytically important in PfuAmyGT),^19^ and E224 (shown to be catalytically important in this paper), find no mention as potentially catalytically-important residues in the papers by Imamura and coworkers.^20,21^ Fig. 5 reveals that residue D362 (shown in green) is present upon a loop in domain 2 (shown in grey). Domain-domain motions can clearly be seen to be required, in order to facilitate involvement of this domain 2 loop in glucose excision and transfer, since the acarbose (shown in ochre) is bound inside a tunnel upon domain 1 (shown in blue) on one side of the loop bearing D362, and the maltose (shown in ochre) is bound to a groove upon domain 2 (shown in grey), with involvement of residues from domain 3 (shown in purple) and domain 1 (shown in blue), on the other side of the same loop. Since the distance between the ends of the acarbose and the maltose happens to be considerably high (> 15 Å), it is necessary either for the loop bearing D362 to move, or for domains 1 and 3 to move with respect to domain 2, for the cleaved glucose unit to be transferred from the end of the tunnel-bound acarbose (analogous to maltotetraose) to the end of a groove-bound maltose.

It is also conceivable that domain-domain motions coupled with subunit-subunit motions occur, because the crystal structure of the *T. litoralis* homolog of PfuAmyGT shows the acarbose and the maltose bound to different subunits of the homodimer. Since there appears to be no route through the subunit interface for a glucose being transferred, it is imperative for the two subunits to be both capable of binding to both of the saccharides. However, it is possible that the catalytic cycle involves an alternation of function between the two subunits of the homodimer, such that motions of the enzyme that cause an acceptor to bind in proximity to the donor in one subunit induce the opposite sequence of binding in the other subunit, i.e., the binding of a donor in proximity to the acceptor, and so on. Indeed, this could explain why (in the *T. litoralis* homolog of PfuAmyGT) Imamura and coworkers found acarbose bound to one subunit, and maltose to the other subunit. The catalytic transfers occurring in the two subunits could thus involve an alternation of the sequence of binding of saccharides between the two binding sites on each subunit, with only one subunit performing catalysis at a time (and driving motions in the other subunit, as a consequence). An alternating and reciprocal mode of action between two subunits could thus potentially explain the working of the enzyme, with the occasional release of a cleaved glucose explaining the appearance of glucose amongst the pool of oligosaccharides that is created each time, regardless of whether the donor is starch, or maltotriose (or a longer malto-oligosaccharide).

Clearly more work will be needed to understand this enzyme completely, and to understand why it possesses three domains, two subunits, a saccharide-binding tunnel as well as a saccharide-binding groove, in addition to four catalytic aspartate/glutamate residues. We candidly admit that despite our identification of four apparently catalytically-important glutamates/aspartates, and our predictions of their pKa values, we have not been able to propose an exact or detailed mechanism of the excision and the transfer. Rather, what we have been able to do very effectively, both in this paper and in the companion paper (using domain-swapped chimeras of PfuAmyGT and a homolog called TonAmyGT from *Thermococcus onnurenius*,^24^ showing that domains 2 and 3 to be involved in determining substrate specificity) is to successfully outlay a detailed challenge to the paradigmatic assumption that there is invariably only a single site and two catalytic glutamates/aspartates in a glucanotransferase, essentially by showing evidence that is incompatible with the operation of single site (while being compatible with separate donor and acceptor binding sub-sites), and demonstrating that there are four catalytic glutamates/aspartates.

## Supporting information

All supplementary tables and figures compiled

## Acknowledgements

We acknowledge funding for this work from three sources: (1) A grant for a Centre for Protein Science, Design and Engineering (MHRD-14-0064) from the Ministry of Human Resource Development (MHRD), Government of India, (2) A project grant for a Hyperthermophile Enzyme Hydrolase Research Centre from the Department of Biotechnology (DBT), Government of India, and (3) Intramural funding from the Indian Institute of Science Education and Research (IISER) Mohali. We also gratefully acknowledge discussions with Dr. Sanjeev Chandrayan and Dr. Neeraj Dhaunta who originally cloned the gene encoding PfuAmyGT in the Guptasarma lab; Dr. Arpana Kumari who carried out some studies with the enzyme’s stability to organic solvents (not reported here); Dr. Nitin Kishore, a past member of the lab, for discussions regarding the mass spectrometry, and Dr. Prajwal Nandekar (Schrödinger) for help with the determination of pKa values.

## Author Credit Statement

AS and PK participated in the conception and execution of experiments performed for this work, and also in all of the analyses of data and the writing up of the manuscript, with PG. PT participated in the early planning and execution, as well as data analysis, with PK. PG supervised the design and execution of experiments, the analyses of data and wrote the manuscript along with AS and PK.

## Accession IDs of protein(s)

The protein referred to as PfuAmyGT has the UniProt Accession No. P49067. It is encoded by the locus no PF0272 in the *Pyrococcus furiosus* in GeneProt. The protein referred to a TonAmyGT from *Thermococcus onnurenius* has the UniProt Accession No. B6YUX8, and TLGT from *Thermococcus litoralis* has the UniProt Accession No. O32462.

## Notes

### Competing Interest Statement

The authors have declared no competing interest.

