## Supplementary material for "A glucanotransferase that uses two sub-sites and four catalytic aspartates/glutamates to disproportionate oligosaccharides ranging in length from maltotriose to starch": All supplementary tables and figures compiled

**Table ST1.**

| Mutants | Primer names | Sequences |
|---|---|---|
| PfuAmyGT wild type | PfuWT_F | 5’-AATAATCATATGATCAATGGTTGGACC-3’ |
| | PfuWT_R | 5’-AATAATCTCGAGACCAGACGCTTC-3’ |
| E32A | E32A_F | 5’-GCGGAAGCGTACGAAAAATGCTACTGG-3’ |
| | E32A_R | 5’-TCGTACGCTTCCGCGAAAACCCAACCG-3’ |
| E33A | E33A_F | 5’-GCGGCGTACGAGAAATGCTACTGGCC-3’ |
| | E33A_R | 5’-CATTTCTCGTACGCCGCTTCGAAAACCCAACCG-3’ |
| E131A | E131A_F | 5’-TGTTTGGCTGACCGCACGTGTTTGGCAG-3’ |
| | E131A_R | 5’-CTGCCAAACACGTGCGGTCAGCCAAACAC-3’ |
| D222A | D222A_F | 5’-GCTGGTGAGAAATTCGGTATCTG-3’ |
| | D222A_R | 5’-GATACCGAATTTCTCACCAGCGTCGTGGAAAACCG-3’ |
| E224A | E224A_F | 5’-GCTAAATTCGGTATCTGGCC-3’ |
| | E224A_R | 5’-CCAGATACCGAATTTAGCACCGTCGTCGTGGAAAAC-3’ |
| E623A | E623A_F | 5’-GCGTCTGGTTGGGACCTGATC-3’ |
| | E623A_R | 5’-GTCCCAACCAGACGCAGACTGAGACAGGG-3’ |

**Table ST1.**

List of primers for achieving the mutant enzymes E32A, E33A, E131A, D222A, E224A and E623A

**Table ST2.**

| mass | position | peptide sequence |
|---|---|---|
| 3993.8592 | 400-434 | DIDYDGFEEVLIENDNFYAV FKPSYGGSLVEFSSK |
| 3106.5209 | 12-37 | INFIFGIHNHQPLGNFGWVF EEAYEK |
| 2948.5628 | 54-78 | VAIHTSGPLIEWLQDNRPEY IDLLR |
| 2862.3396 | 165-188 | EELYWPYYTEDGGEVIAVFP IDEK |
| 2760.2688 | 448-470 | WEHYHGYVESQFDGVASIHE LEK |
| 2740.4701 | 561-586 | TGNPVLFAVELNVAVQSIME SPGVLR |
| 2576.3354 | 618-640 | TLSQSESGWDLIQQGVSYIV PIR |
| 2483.2097 | 488-508 | FMLQDHVVPLGTTLEDFMFS R |
| 2417.1607 | 358-378 | AQCNDAYWHGLFGGVYLPHL R |
| 2401.2148 | 271-291 | GLVYLPIASYFEMSEWSLPA K |
| 2346.0958 | 144-164 | ESGIDYVIVDDYHFMSAGLS K |
| 2116.1688 | 84-103 | GQVEIVVAGFYEPVLASIPK |
| 2062.9289 | 38-53 | CYWPFLETLEEYPNMK |
| 1774.8475 | 226-239 | FGIWPGTYEWVYEK |
| 1752.8802 | 546-560 | LVNDGFEVEYIVNNK |
| 1717.8643 | 201-215 | VLEYLHSLIDGDESK |
| 1529.7920 | 255-266 | INLMLYTEYLEK |
| 1481.7634 | 518-530 | VPYSYELLDGGIR |
| 1249.5881 | 1-11 | MINGWTEVGDK |
| 1247.7146 | 191-200 | YLIPFRPVDK |
| 1237.6827 | 295-304 | LFVEFVNELK |
| 1212.5241 | 605-613 | FEDEMEVWK |
| 1168.5092 | 328-336 | YPESNYMHK |
| 1161.6626 | 437-446 | LVNYVDVLAR |

| mass | position | peptide sequence |
|---|---|---|
| 1103.5480 | 509-517 | QQEIGEFPR |
| 1053.5575 | 534-542 | EHLGIEVEK |
| 998.5669 | 133-140 | VWQPELVK |
| 971.5672 | 380-387 | AIWNNLIK |
| 938.4941 | 388-396 | ANSYVSLGK |
| 860.4624 | 126-132 | GVWLTER |
| 817.4301 | 589-595 | EIVVDDK |
| 765.3889 | 119-125 | SIGFDAR |
| 742.4093 | 472-477 | IPDEIR |
| 724.3512 | 479-484 | EVAYDK |
| 718.3559 | 323-327 | NFFYK |
| 713.3253 | 244-248 | EFFDR |
| 708.3783 | 338-343 | MLMVSK |
| 700.3373 | 347-352 | NNPEAR |
| 678.3304 | 249-254 | ISSDEK |
| 658.3882 | 107-111 | IEQIR |
| 639.2620 | 651-656 | FEEASG |
| 593.3293 | 307-311 | GIFEK |
| 564.3504 | 354-357 | YLLR |
| 563.3300 | 267-270 | YKPR |
| 560.3191 | 318-322 | GGIWK |
| 537.3031 | 596-600 | YAVGK |
| 533.2718 | 115-118 | EWAK |
| 531.3038 | 240-243 | GWLR |
| 520.3242 | 314-317 | VFVR |
| 506.2973 | 614-617 | YPVK |
| 504.2664 | 641-644 | LEDK |

**Table ST2.**

List showing the *in silico* (tryptic) digested peptides of wild type PfuAmyGT.

**Table ST3.**

| **Compound** | **Expected MW of compounds (as sodium adducts) (g/mol)** | **Observed MW of compounds (as sodium adducts)** |
|---|---|---|
| Glucose | 203.1457 | - |
| Maltose | 365.2898 | 365.1023 |
| Maltotriose | 527.4268 | 527.1697 |
| Maltotetraose | 689.5698 | 689.2419 |
| Maltopentaose | 851.7098 | 851.3149 |
| Maltohexaose | 1,013.8487 | 1013.3824 |
| Maltoheptaose | 1,175.9898 | 1175.4396 |

**Table ST3.**

Theoretical and observed masses of glucose, maltose and other malto-oligosaccharide products (as sodium adducts) following hydrolysis of substrates by PfuAmyGT

**Fig. S1.**

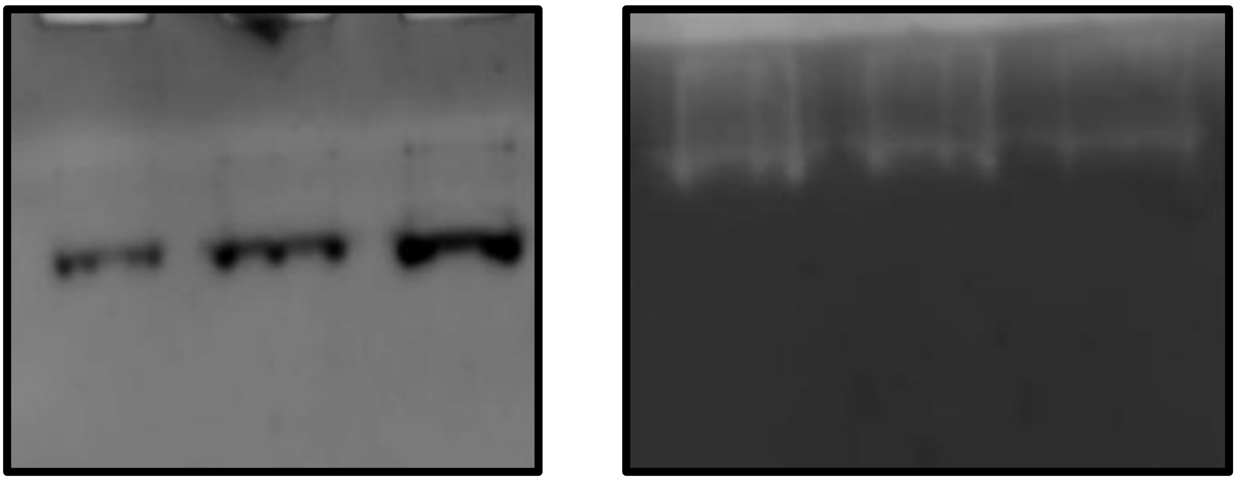

**Fig. S1.**

Native Page and zymogram of PfuAmyGT: The left panel shows bromophenol blue stained native PAGE of full-length Pfu AmyGT which was loaded in different (increasing) concentrations. Right panel shows iodine stained native PAGE gel which shows activity in form of clear zones.

**Fig. S2.**

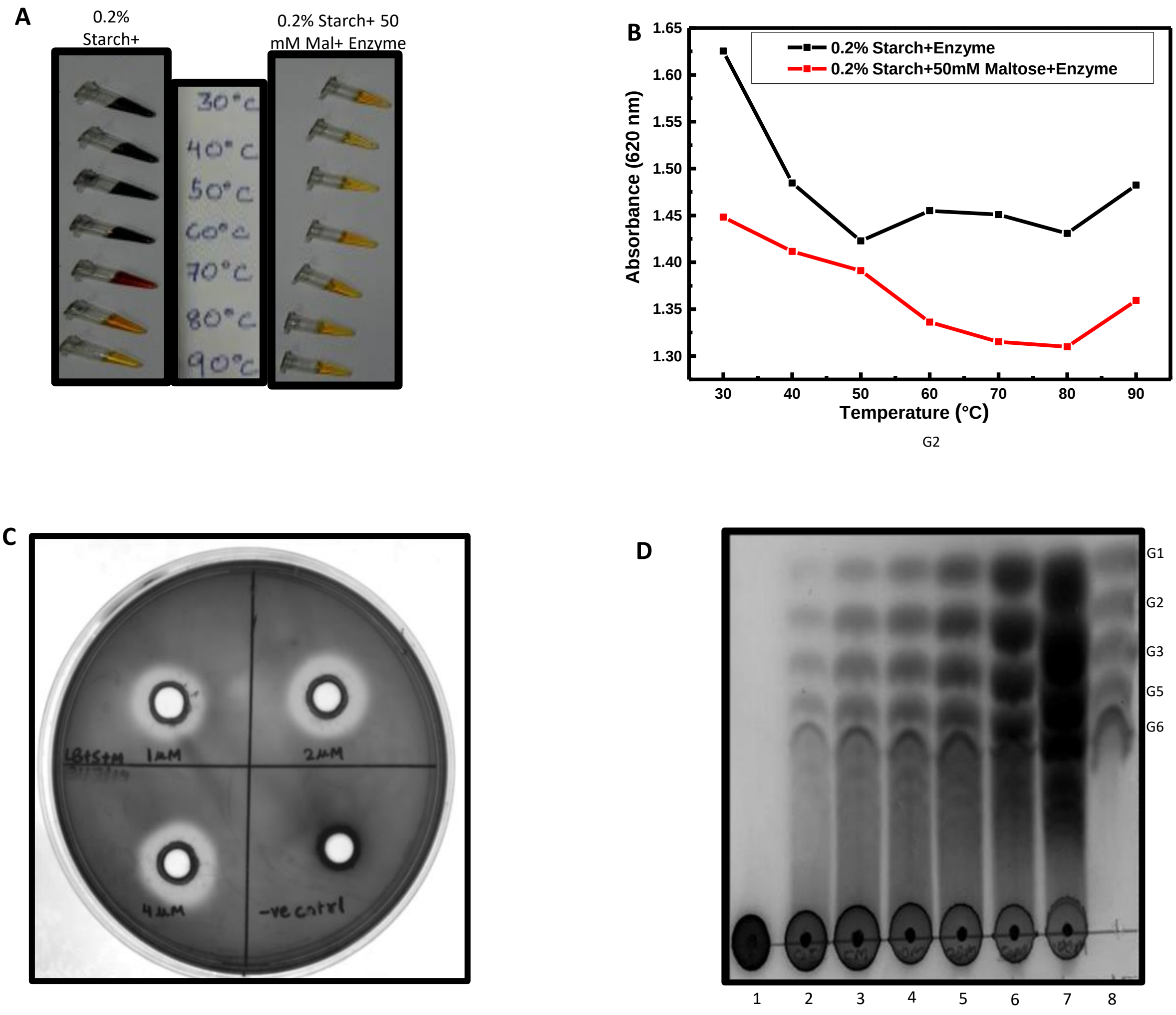

**Fig. S2.**

A. Visual representation of amylase activity obtained by the action of full-length PfuAmyGT on 0.2% (left panel) and 0.2% starch + 50 mM Maltotriose (right panel) at different temperatures (right panel)

B. Detection of amylase activity by the starch-iodide method: Hydrolysis of starch observed as decreased absorbance at 620 nm, in the absence or presence of maltose.

C. Visual representation showing clear zones on an agar plate containing 0.2 % starch and 50 mM maltose and increasing concentration of wild type PfuAmyGT.

D. TLC showing an increasing intensity of oligosaccharide products with PfuAmyGT (1 µM) with increasing maltose concentration (incubation carried out at 90 deg Celcius for 12 hours): **Lane 1** shows the control reaction with 1% starch without the enzyme; **Lane 2** shows the activity profile of the enzyme with 1 % starch; **Lane 3** shows the activity profile of the enzyme with 1 % starch and 5 mM Maltose; **Lane 4** shows the activity profile of the enzyme with 1 % starch and 10 mM Maltose; **Lane 5** shows the activity profile of the enzyme with 1 % starch and 20 mM Maltose; **Lane 6** shows the activity profile of the enzyme with 1 % starch and 50 mM Maltose; **Lane 6** shows the activity profile of the enzyme with 1 % starch and 100 mM Maltose; **Lane 7** shows standard oligosaccharides obtained commercially and pre-mixed (20 mM each) and chromatographed.

**Fig. S3.**

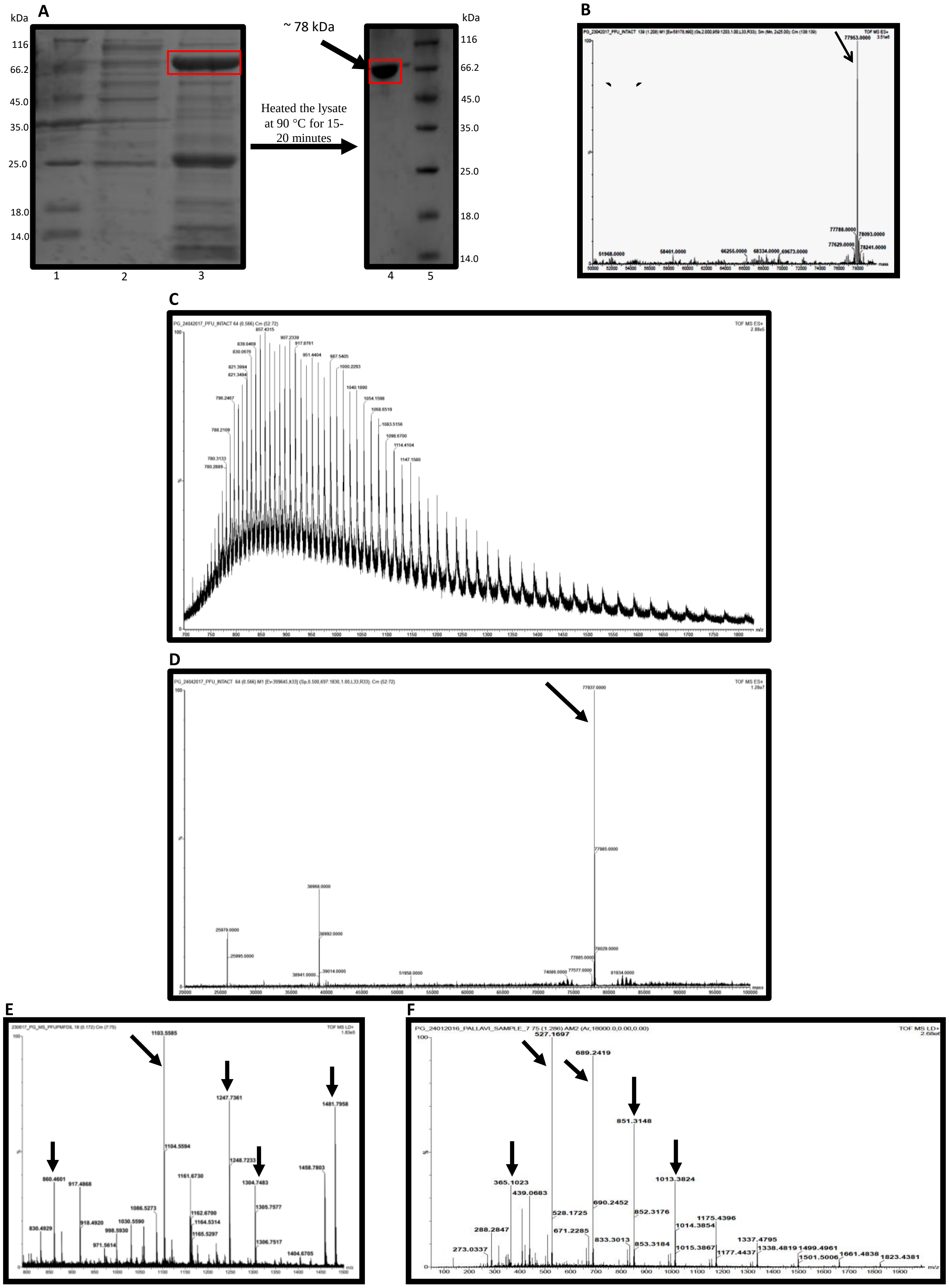

**Fig. S3.**

A. SDS PAGE analysis of PfuAmyGT showing the effect of heat treatment in preventing degradation and improving purification of intact protein.

B. Deconvoluted mass spectrum of intact PfuAmyGT (zoomed to show the mass range of ~50,000-80,000 Da), obtained using ESI-Q-TOF MS.

C. Raw mass spectrum of intact PfuAmyGT (zoomed to show the m/z range of ~700-1900) showing various charged states used to obtain the deconvoluted mass shown in (b) above, obtained using ESI-Q-TOF MS.

D. Full range (~20,00-1,00,000 Da) deconvoluted mass spectrum of intact PfuAmyGT sample showing the dominant mass peak of 77937 Da, along with some other minor contaminating species, obtained using ESI-Q-TOF MS.

E. Peptide mass fingerprint (PMF) mass spectrum showing peptides obtained after tryptic digest of PfuAmyGT (with arrows annotating peptide masses matching expected masses, within 100 ppm accuracy), obtained using MALDI-Q-TOF MS.

F. Mass spectrometric analysis of sugars obtained as hydrolyzed products from the reaction involving 1 % starch + 20 mM maltose + 1 µM PfuAmyGT, shown as a representative mass spectrum of sodium adducts of sugars, obtained using MALDI-Q-TOF MS. Theoretical and

**Fig. S4.**

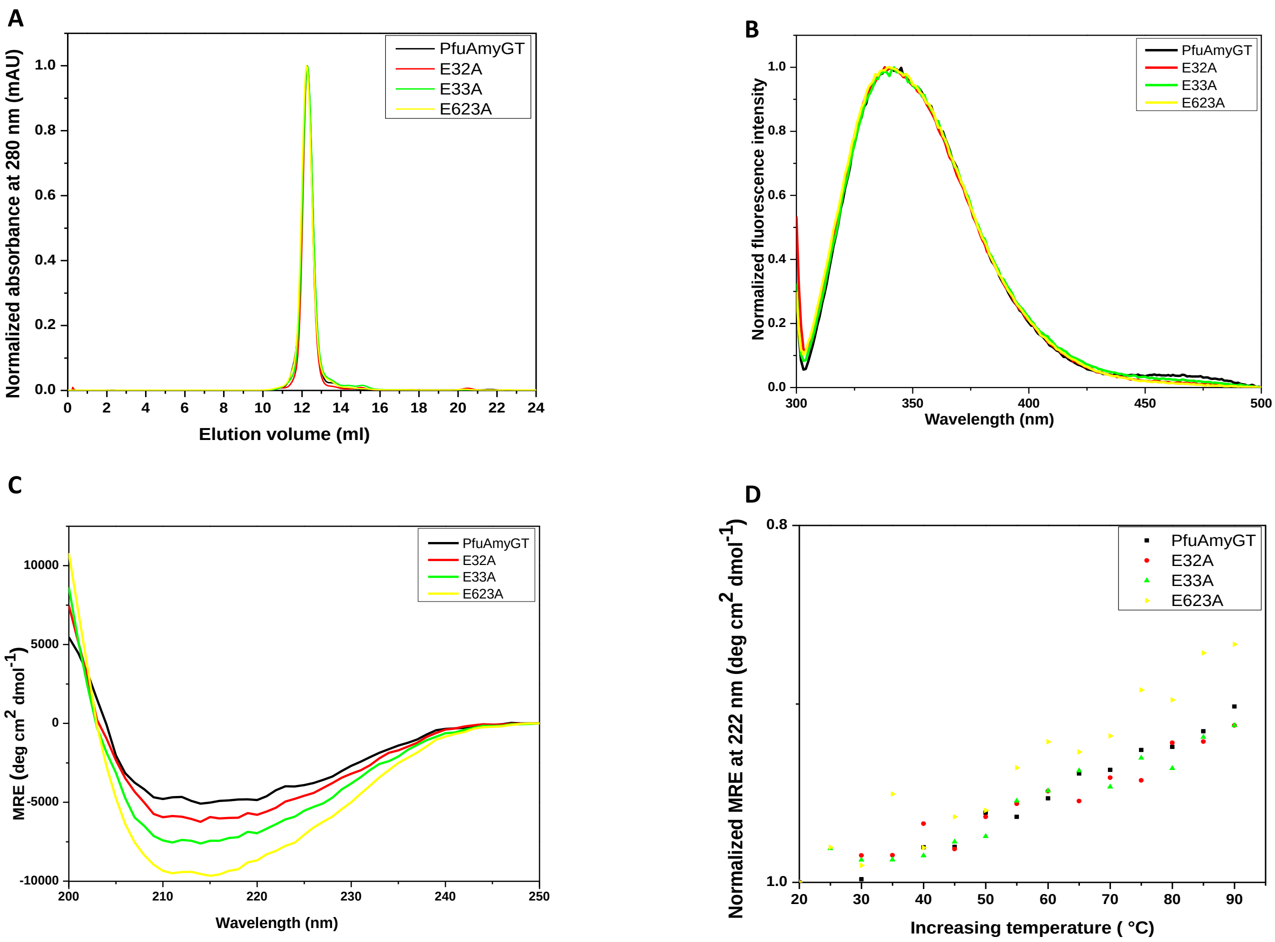

**Fig. S4.**

A. Gel filtration chromatographic analysis of wild type E32A, E33A and E623A.

B. Normalized fluorescence emission spectrum wild type E32A, E33A and E623A.

C. Circular Dichroism (CD) spectrum of wild type E32A, E33A and E623A.

D. Thermal melt at 222 nm of wild type E32A, E33A and E623A.

**Fig. S5.**

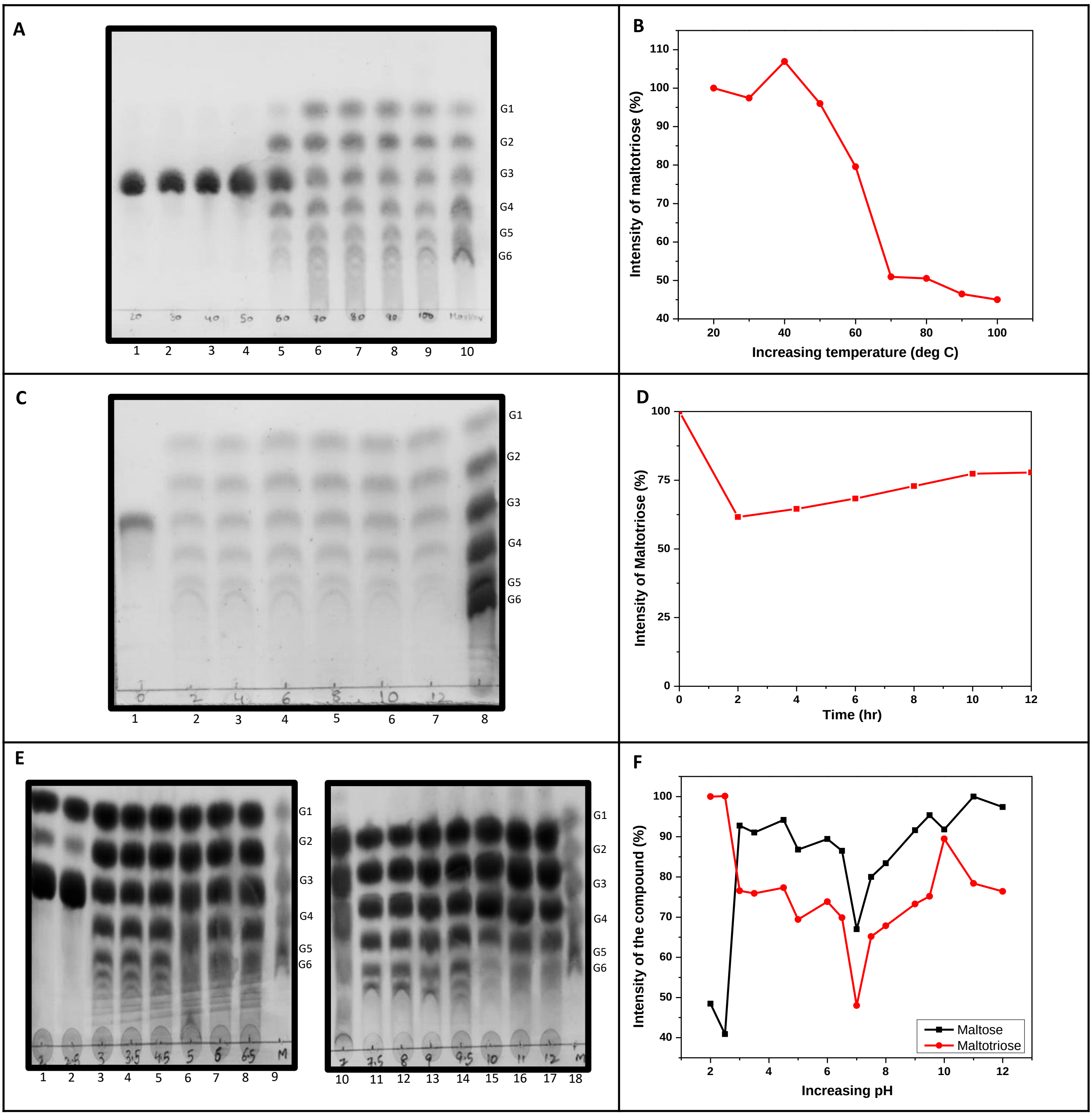

**Fig. S5.**

A. TLCs showing the temperature profile of wild type PfuAmyGT with 20 mM Maltotriose as the sole substrate: **Lane 1** to **Lane 4** shows the reaction products of 1 uM wild type PfuAmyGT with 20 mM of Maltose incubated for 12 hours at 20 deg Celcius, 30 deg Celcius, 40 deg Celcius and 50 deg Celcius, respectively; **Lane 5** shows standard oligosaccharides obtained commercially and pre-mixed (20 mM each) and chromatographed.; **Lane 6** to **Lane 10** the reaction products of 1 uM wild type PfuAmyGT with 20 mM of Maltose incubated for 12 hours at 60 deg Celcius, 70 deg Celcius, 80 deg Celcius and 90 deg Celcius, respectively; **Lane 11** shows the standard oligosaccharides obtained commercially and pre-mixed (20 mM each) and chromatographed.

B. The graph represents the change in intensity of maltotriose (in percentage) as a function of temperature.

C. Time-dependent profile of wild type PfuAmyGT activity with 20 mM Maltotriose as the sole substrate: **Lane 1** shows the control reaction with 20 mM Maltotriose without the enzyme; **Lane 2** to **Lane 7** shows the reaction products of 1 uM wild type PfuAmyGT with 20 mM of Maltotriose incubated at 90 deg Celcius for 2 hours, 4 hours, 6 hours, 8 hours, 10 hours and 12 hour respectively; **Lane 8** shows the standard oligosaccharides obtained commercially and pre-mixed (20 mM each) and chromatographed.

D. The graph represents the change in intensity of maltotriose (in percentage) as a function of time.

E. pH-dependent profile of wild type PfuAmyGT activity with 20 mM Glucose and 20 mM Maltotriose : **Lane 1** to **Lane 8** shows the reaction products of 1 uM wild type PfuAmyGT with 20 mM Glucose and 20 mM of Maltose (incubated at 90 deg Celcius for 12 hours) at pH 2, 2.5, 3, 3.5, 4.5, 5, 6 and 6.5 respectively. **Lane 9** shows the standard oligosaccharides obtained commercially and pre-mixed (20 mM each) and chromatographed; **Lane 10** to **Lane 17** shows the reaction products of 1 uM wild type PfuAmyGT with 20 mM Glucose and 20 mM of Maltose (incubated at 90 deg Celcius for 12 hours) at pH 7, 7.5, 8, 9, 9.5, 10, 11 and 12 respectively; **Lane 18** shows the standard oligosaccharides obtained commercially and pre-mixed (20 mM each) and chromatographed.

F. The graph represents the change in intensity of maltose and maltotriose (in percentage) as a function of pH.
